# PHYFUM: Phylogenetic Reconstruction of Normal and Pre-malignant Tissue Evolution Using Fluctuating Methylation

**DOI:** 10.64898/2026.05.05.722981

**Authors:** Pablo Bousquets-Muñoz, Heather E. Grant, Darryl Shibata, Trevor A. Graham, Carlo C. Maley, Calum Gabbutt, Diego Mallo

**Author notes:** Co-senior authors (Calum Gabbutt, Diego Mallo).

## Abstract

We present PHYFUM, a novel Bayesian phylogenetic method for methylation data that reconstructs the evolutionary history of stem cells and the glandular structures they reside within in normal tissue. Using simulations, we validated this phylogenetic method and confirmed its accuracy. A re-analysis of 22 patients unveiled early gland divergence in the human gut, in contrast to a much later common ancestor in the endometrium, and yielded strong evidence against gland division by segregation of individual stem cells.

## Main

Cancer development and normal tissue homeostasis are dynamic evolutionary processes that are difficult to observe *in vivo* in humans. Population genetic and phylogenetic models leverage heritable alterations to reconstruct evolutionary relationships between cells. Many existing methods are constrained by the relatively low nucleotide mutation rate, and most studies are limited by the number of samples and patients, in part due to the high cost of deep whole-genome sequencing. Here, we present a novel probabilistic phylogenetic method that leverages somatic DNA methylation alterations to trace clonal evolution at low cost and unprecedented temporal resolution.

Within glandular epithelium, such as intestinal and endometrial glands, stem cells reside in a niche and undergo neutral competition, with stem cells repeatedly replacing their neighbours^1^. We recently discovered fCpGs, CpG dinucleotide sites that undergo stochastic methylation fluctuations^2^. Our accompanying *Flip-Flop* method fits a mathematical model to the characteristic W-shaped methylation fraction distributions, allowing us to estimate the dynamics of a single stem-cell niche. This method was able to estimate critical homeostatic parameters, such as the number of stem cells (niche size), stem-cell replacement rate, and (de)methylation rates^1^ (hereafter stem-cell-niche parameters), but was agnostic to the inter-gland evolutionary relationships and changes in their evolutionary dynamics over time and space. However, during development and adult life, intestinal glands can bifurcate and clonally expand in a process known as crypt fission^3^. Here, we present *PHYFUM* (PHYlogeny from FlUctuating Methylation), a phylogenetic application of our *Flip-Flop* model^2^ that enables the simultaneous inference of these stem-cell-niche parameters and the evolutionary relationships between glands (Fig. 1). We previously developed a simple phylogenetic approach that collapsed each clonal population to the methylation state of its inferred most recent common ancestor (MRCA), enabling reconstruction of relationships between bulk samples but ignoring intra-population dynamics^4^. In contrast, PHYFUM draws upon polymorphism-aware species-tree phylogenetic methods^5^ by explicitly modelling methylation fluctuations and stem-cell replacement directly along phylogenetic branches, while the tree tracks the divergence between glands.

**Fig. 1:**
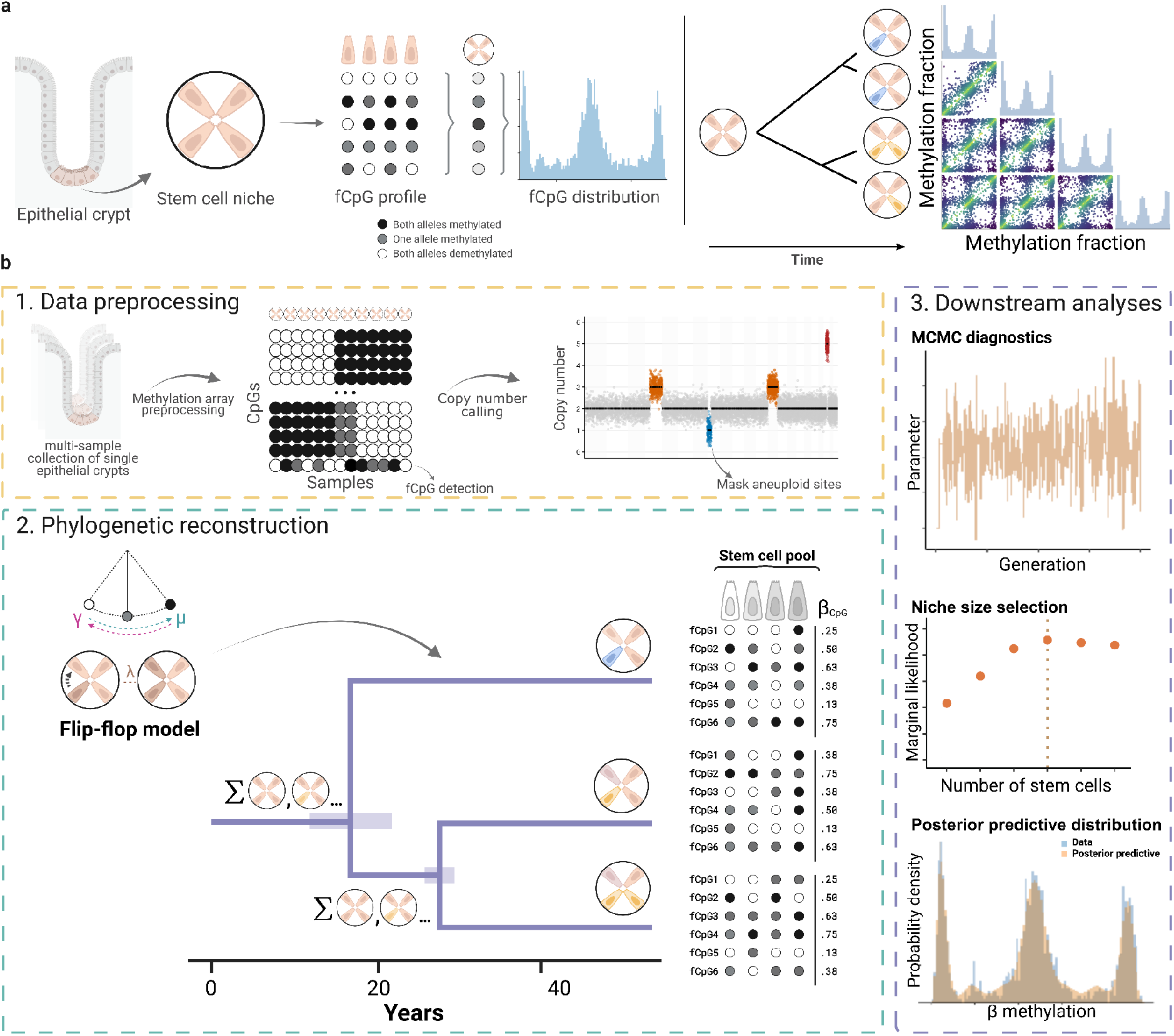
Fluctuating methylation enables phylogenetics of normal epithelial tissues. **a**, Bulk methylation of oligoclonal populations (e.g., epithelial glands) at fluctuating CpG sites reflects the methylation status of the population stem cells. As glands divide, fCpG sites randomly diverge, and pairwise differences can be observed across samples, albeit the stable W-shaped distribution of fCpG per sample. **b**, PHYFUMflow seamlessly performs the preprocessing of raw data (1), phylogenetic reconstruction with PHYFUM (2), and downstream analyses with PHYFUMr involving QC, stem-cell-niche size comparison with selection or averaging, and plots, among others (3).

Built upon the *BEAST* phylogenetic framework^6^ and our previous adaptation to study somatic evolution *PISCA*^7^, PHYFUM implements substitution matrices and data-uncertainty models adapted from our previous fCpG method, new gland division models, and modifications to the pruning algorithm^7^ needed to study somatic evolution (Fig. 1a). Intra-niche evolution is modeled as a non-reversible continuous-time Markov Chain. The discrete states in this chain represent the niche’s methylation status (the number of fully methylated, half-methylated, and unmethylated cells) at each fCPG site, which transition via methylation, demethylation, or cell replacement (see Gabbutt et al.^2^, and online methods). To map the observed methylation fractions to these discrete states, we employ a model that accounts for measurement error. Inter-niche evolution is described by the phylogenetic tree and is compatible with standard demography models (e.g., coalescent). Internal nodes represent ancestral stem-cell niches that, by default, are assumed to generate two identical daughter niches when they bifurcate. The phylogenetic likelihood is calculated using the Felsenstein pruning algorithm^8^, thus marginalising over all possible ancestral states of the internal nodes. The tree is rooted at an additional external node at the origin of the process (i.e., patient birth) with an assigned prior state frequency distribution. The size of the stem-cell niche is estimated using Bayesian model comparison or marginalised *a posteriori* using Bayesian model averaging.

We developed a phylogenetic fCpG simulator and used it to validate PHYFUM’s performance. Within a realistic parameter space that is consistent with our previous work^2^ PHYFUM showed exceptional accuracy, with a median relative error (|Posterior mean - Expected|/Expected) of 0.056 (IQR: 0.019) across all continuous parameters (n conditions = 50 with 3 replicates, Fig. 2a). Tree reconstruction was similarly accurate, with a median weighted Robinson-Foulds distance^9^ to the true tree of 0.038 (IQR: 0.045) (0≤wRF≤1, considering topology and branch lengths). Our previously published single-cell phylogenetic method was significantly less accurate, showing a 0.14 (IQR: 0.14) median tree error (Fig. 2b; paired Wilcoxon signed-rank test, p < 0.001). Using a much broader simulation space (n = 486; Supplementary Figure 1), we identified exceptional parameter values where PHYFUM underperforms due to imbalanced or very fast epigenetic switching rates (≳10^-1^ fluctuations per year in typical adult samples). Crucially, both exceptions can be detected in the input data. Imbalanced rates do not generate the characteristic W-shape fCpG distributions (Supplementary Figure 2) and are discarded during fCpG selection. Fast rates lead to phylogenetic saturation, which is detected by two new statistics included in our software (see Supplementary Methods; validated in a second simulation study [n = 42, Supplementary Figures 3-4]). Large stem-cell niches (*S*) are more difficult to estimate. Estimation under a smaller *S* yields a minor error penalty in parameters unrelated to *S* and may be an adequate compromise when estimating *S* is not a primary concern (Supplementary Figures 5-6).

**Fig. 2:**
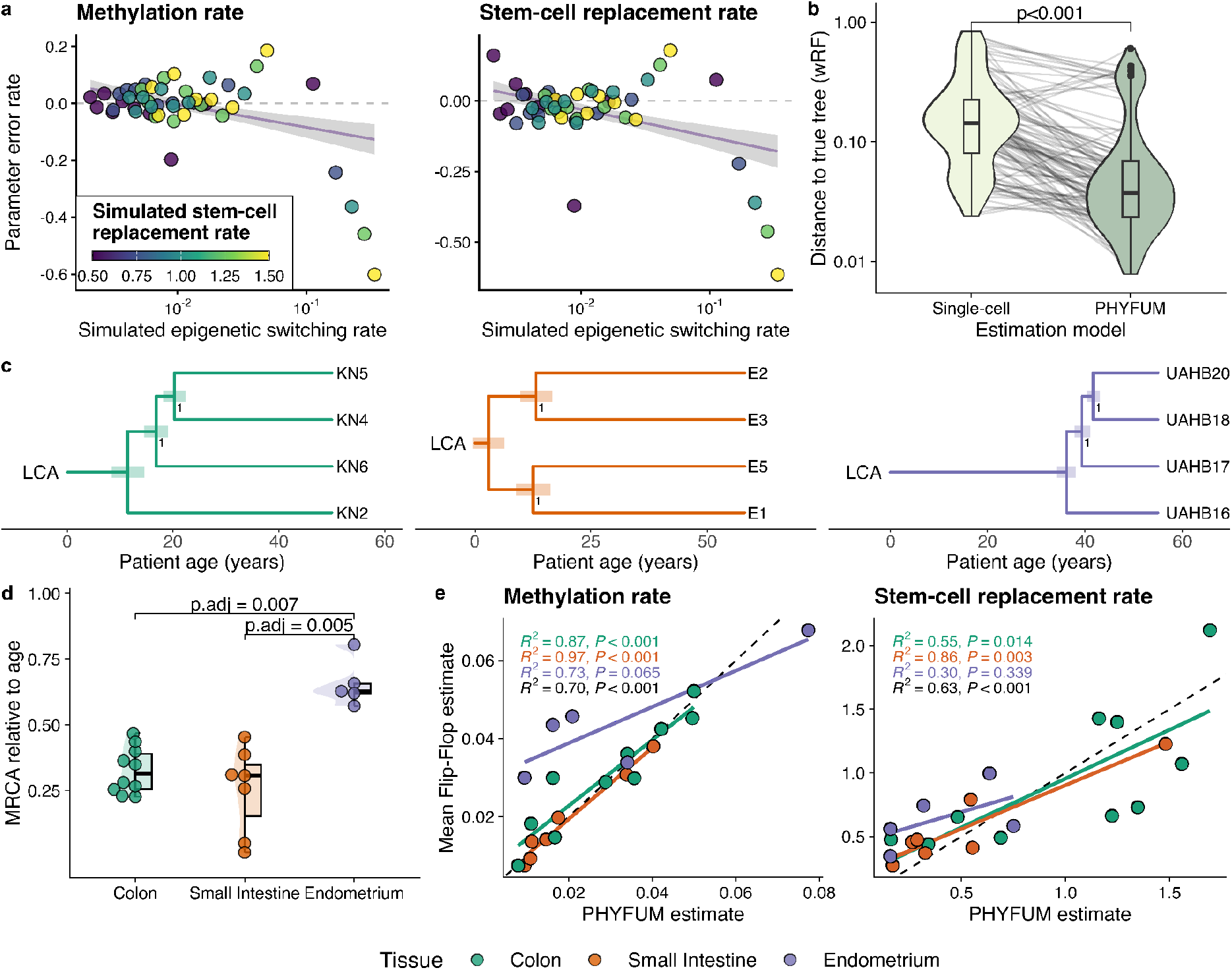
Performance of PHYFUM on simulated and real data. **a**, Error rate on substitution model parameters in a range of simulated rates of epigenetic switching (sum of methylation and demethylation rate) and stem cell replacement. Demethylation rate error is highly correlated with methylation rate. Robust linear regression models the data with outliers, which reveal the model’s accuracy limits. **b**, Comparison of inferred trees distance to the ground truth (weighted Robinson Foulds distance) against single-cell simplification of PHYFUM’s model and PHYFUM. **c**, Representative phylogenetic trees in the three studied tissues (colon, small intestine, and endometrium). **d**, Proportion of the evolutionary time taken by the LCA-MRCA branch. Dunn test with Holm-adjusted p-values for significant contrasts. **e**, Comparison of real-data parameter estimates between PHYFUM and *Flip-Flop*^2^. Single-sample estimates calculated with the *Flip-Flop* model were averaged by patient for comparison.

We next applied PHYFUM to a set of bulk human healthy epithelial glands from the colon, small intestine, and endometrium, collected from each donor sample at approximately 1 cm intervals. The inferred evolutionary histories of the three tissues differed markedly (Figure 2c, Supplementary Figure 7). Endometrium samples diverged more recently than gut samples, which was reflected in the proportion of relative evolutionary time happening before the MRCA of the samples (Fig.2d, adjusted p-values < 0.01), and their absolute timing (colon: 15.1 ± 4.1 years [mean ± 95% CI]; intestine: 12.4 ± 4.9; endometrium: 29.9 ± 5.7). This likely represents true underlying biology related to endometrial glandular replacement during menstrual cycles. In contrast, the intestine presented phylogenies with the MRCA shortly after birth, branching during puberty, and long terminal branches extending into adulthood, consistent with normal gland division during development^3,10,11^. Stem-cell niche parameters were similar between colon and small intestine, but different in endometrium (Supplementary Table 1). The switching rate was faster in endometrial tissue compared to colon (p.adj_demethylation rate_ <0.001, p.adj_methylation rate_ = 0.07), and the stem-cell replacement rate was slower (p.adj=0.03). All stem-cell-niche model parameters estimates showed high consistency between PHYFUM and *Flip-Flop* (median Pearson correlation r = 0.77, each with p < 0.001), except for the number of stem cells (r = 0.17, p = 0.4, Fig.2e, Supplementary Figure 8-9). This discordance in estimates of the number of stem cells is largely due to differences in the parameterization of their error models. Using the same error model, the correlation is present and of similar strength as that of the other parameters (r = 0.5, p < 0.001; Supplementary Figure 10). When stratified by tissue, concordance between PHYFUM and *Flip-Flop* estimates was weakest in the endometrium. This likely reflects endometrium-specific estimation instability in *Flip-Flop*, as the replacement rate estimates were decoupled from the epigenetic switching rates exclusively in endometrium (Supplementary Figure 11). PHYFUM does not show this pattern, suggesting that jointly modelling multiple samples allows it to overcome this parameter identifiability issue.

There is ongoing debate about how glands divide in healthy and pre-malignant tissues, with the classical view of each daughter gland receiving an equal number of stem cells (symmetric gland fission) being challenged by murine studies indicating a budding process in normal colon^12^ and human pathology studies reporting asymmetrical branching in colorectal adenomas^13^. PHYFUM models stem-cell-niche division at each internal node, enabling comparisons among models. In addition to our default model above (which assumes stem-cell niches are identical after gland-division, hereafter called *Cloning*), we implemented three asymmetric gland-division models (Fig. 3A): *Double-Split*, in which the stem cells first double and then are randomly assigned to the two daughter glands; *Split-Double*, in which a random split of the stem cells occurs first and then they double; and *Budding*, in which a single cell divides and one of its daughters buds from the niche, establishing a new niche *de novo* (see Online Methods).

**Fig. 3:**
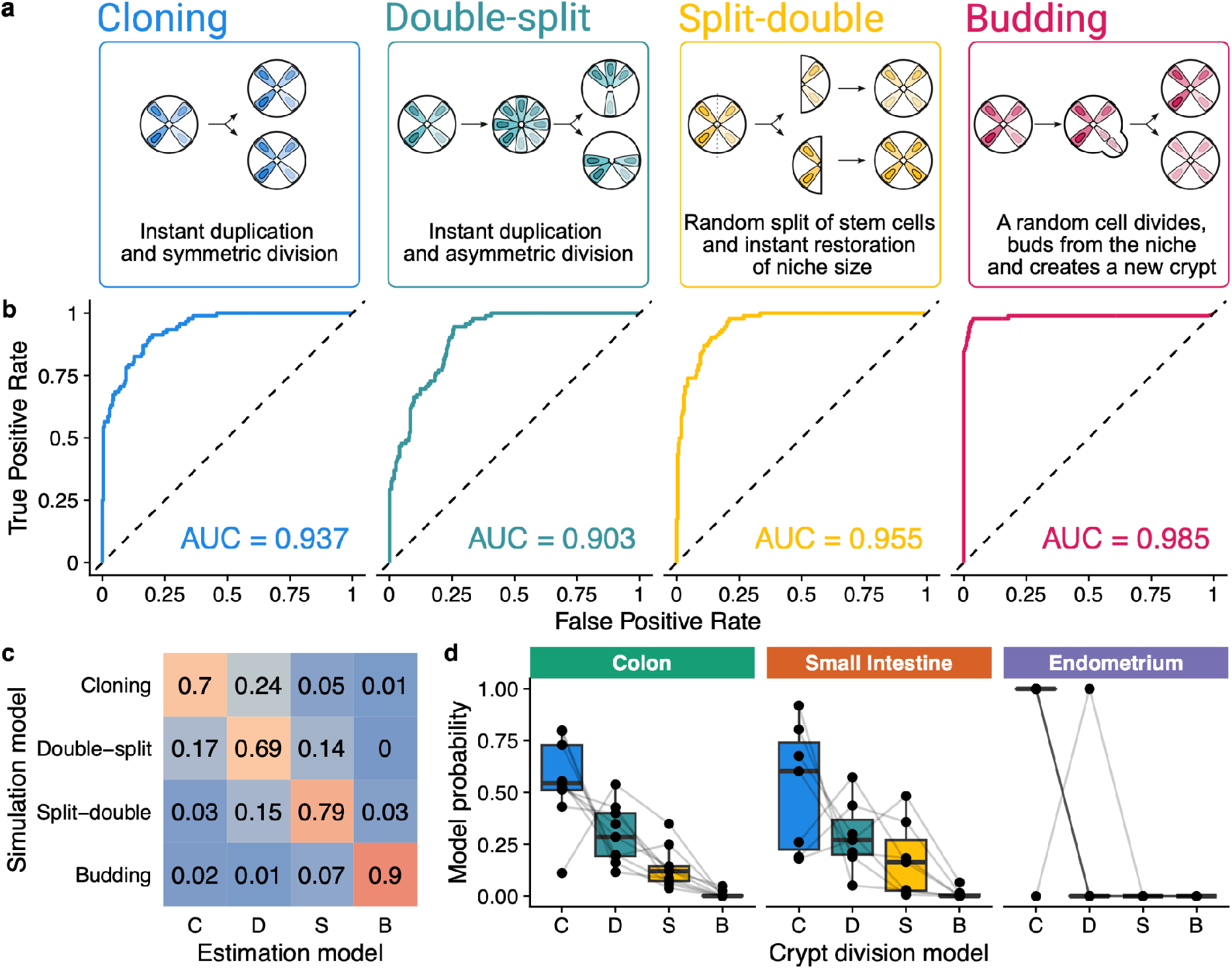
Gland-division model selection with PHYFUM. **a**, Schematics of the four gland-division models implemented in PHYFUM. **b**, Multiclass ROC curves of the model of interest vs. the rest showing PHYFUM’s gland-division classification performance in simulations. **c**, Confusion matrix indicating the proportion of simulations for which the gland-division model was estimated correctly (diagonal) and incorrectly (off-diagonal). **d**, Distribution of gland-division model posterior probabilities per patient stratified by tissue. Model abbreviations: C=Cloning, D=Double-split, S=Split-double, B=Budding.

We repeated the previous 150 simulations under the four models and performed phylogenetic estimation for each under each model, generating all 16 combinations to study model identifiability via Bayesian model selection and the effect of model misspecification on PHYFUM’s accuracy. There was excellent classification performance across the board (AUCs >0.9 for all models indicated, Fig. 3B), with *Budding* being the most distinguishable. The confusion matrix (Fig. 3C) confirmed these findings and showed that the models that generate larger state changes at gland-division (*Budding* and *Split-Double*) are easier to distinguish, whereas *Double-Split* and *Cloning* are more similar, leading to a higher misclassification rate between them. This was expected, given that *Double-Split* is the only alternative model that allows the same stem cell lineage to seed both daughter glands. Model distinguishability decayed with the evolutionary distance to the last internal node. This is illustrated by the strong correlation between the probability of the *Budding* model in simulations with other models and the evolutionary distance from the sampling time to the most recent internal node (Spearman’s rank correlation ρ = 0.78, p < 0.001). Model misspecification induced moderate estimation error, peaking at a 66% increase in median tree reconstruction error (*Double-split* simulations estimated using *Budding*). The true model always showed the lowest error rate, and the incorrect alternatives significantly increased it (Supplementary Figures 12-13). When the inference model was misspecified, *Budding* consistently yielded the largest reconstruction errors, whereas *Double-split* was generally the most robust.

We used PHYFUM to compare gland fission dynamics in healthy human intestine and endometrium (Supplementary Figure 14a). *Cloning* best fitted the data in most cases (17/22), with weak evidence against the other asymmetric fission models in intestinal samples. Interestingly, model distinguishability was dramatically higher in endometrial samples. This finding is at least partially explained by its much more recent internal nodes meaning stem cells had closer relationships between glands (Fig. 2d-e), in line with our observation in simulated data. *Budding* was strongly rejected across tissues (Fig. 3D, Supplementary Figure 14b). This finding, in combination with our model’s high *in silico* identifiability, constitutes strong evidence against budding as the mechanism of gland-division in our samples.

In summary, PHYFUM is a novel tool that accurately reconstructs phylogenies from healthy oligoclonal samples by leveraging fluctuating methylation. Input data can be obtained using cost-effective arrays, making this approach practical for large-scale studies. Here, PHYFUM revealed distinct gland evolution patterns between the intestine (colon and small intestine) and the endometrium, and provided strong evidence against gland division by budding in these healthy human tissues. This highlights its potential to enhance our understanding of healthy tissue dynamics and to develop evolutionary biomarkers of neoplastic progression.

## Software Availability

All the software developed for this manuscript is distributed under open-source licenses. The core phylogenetic method, PHYFUM, is available at https://github.com/pbousquets/phyfum. To ensure broad accessibility and ease of use, we developed two accompanying pieces of software. PHYFUMr (https://github.com/adamallo/PHYFUMr) is an R package that enables users to manually prepare input data and analyze PHYFUM’s results, including data preprocessing, copy number variant detection to exclude aneuploid CpG sites, *de novo* identification of fCpG sites, phylogenetic analysis, Bayesian model comparison and/or model averaging for S and gland-division model, quality control, summarization, and publication-ready figure generation. PHYFUMflow (https://github.com/pbousquets/phyfumflow) is an end-to-end Snakemake workflow that automates the execution of PHYFUM and PHYFUMr. It is distributed as a Python package and as a Docker image bundled with PHYFUM, providing a streamlined solution for researchers studying clonal dynamics in healthy and neoplastic tissues. The complete suite of tools is, and will always be, open-source. Code to replicate this study is available at https://github.com/adamallo/PHYFUM_manuscript.

## Acknowledgments

This research was primarily funded by the US National Institutes of Health National Cancer Institute (U54 CA217376 to C.C.M., D.S., and T.A.G., R01 CA140657 and R01 CA285517 to C.C.M.), Cancer Research UK (EDDPMA-May23/100059 to T.A.G. and C.G. and DRCNPG-May21_100001 to T.A.G.), ARPA-H (ADAPT 140D042590007 TO C.C.M.), the Colorectal Cancer Alliance (to C.C.M.), and the Thornton Foundation (to T.A.G. and C.G.). C.G. acknowledges support from the Chapman-Schmidt AI in Science scheme at Imperial College London. Boehringer Ingelheim Fonds provided funding to support P.B. in this research project.

The authors acknowledge Research Computing at Arizona State University for providing storage and computation resources on the Sol supercomputer, which were essential to the research results reported in this paper.

## Author Contributions

C.G., T.A.G., D.M., and C.C.M. conceived the project. C.G., D.S., and T.A.G. developed the original theoretical framework. D.M., C.G., P.B., and H.E.G. designed the study and phylogenetic methods. C.G. designed, implemented, and conducted non-phylogenetic methods and analyses, and developed and implemented the simulator. P.B. and D.M. implemented the phylogenetic methods and conducted the phylogenetic analyses with H.E.G.. P.B. and D.M. drafted the manuscript and figures. T.A.G., C.C.M., P.B., C.G. and D.S. secured funding for the project. All authors interpreted the results, revised the manuscript critically for important intellectual content, read, and approved the final manuscript.

## Online Methods

### 1. PHYFUM: a phylogenetic model of fluctuating methylation

BEAST provides a framework for Bayesian phylogenetic reconstruction with an extensive collection of models that typically utilise substitutions (i.e., single-nucleotide replacements) within species or populations. PHYFUM builds on BEAST 1.8^6^ to introduce models for fluctuating methylation in non-tumoral samples of cell populations originating from a single stem-cell niche (e.g., single glands). It integrates three primary new components:

- **Stem-cell niche model:** describes the evolution within a stem-cell niche as a continuous-time Markov Chain given by 4 parameters: methylation rate (*μ*), demethylation rate (*γ*), stem-cell replacement rate (λ), and the number of stem cells in the niche (*S*).
- **Error model:** maps the continuous (proportion) data to the discrete evolutionary states probabilistically, modeling methylation array intensities by accounting for background signal and compression near the assay bounds (ϵ and Δ), and the assay’s precision *κ*.
- **Tree model:** describes the evolutionary relationships between stem-cell niches and the time to their ancestors. It requires modifications to the phylogenetic likelihood calculation and includes different models of how glands divide at internal nodes.

Additionally, the demographic model, which uses the tree shape to estimate other evolutionary dynamics between stem-cell niches, and the clock models, which estimate rate variation over time, have not been meaningfully modified in PHYFUM.

Two PHYFUM parameters, the number of stem-cells in the niche *S* and the gland-division model, are not estimated within the MCMC and therefore require independent runs and subsequent Bayesian model comparison or marginalisation.

More details on ancillary implementations needed to integrate these models with BEAST can be found in the Supplementary Methods.

#### 1.1. Fluctuating methylation model

The continuous-time Markov Chain model that describes the evolution of a single stem cell niche is identical to the original *Flip-Flop* implementation^2^. In brief, for a single fCpG, a stem cell niche of fixed size *S* transitions between all possible 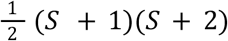 combinations of each of those cells being fully methylated *m*, half methylated *k*, or unmethylated *S* − *k* − *m*. Transitions are explained by three mutually exclusive events with independent rates: methylation *μ*, demethylation *γ*, and cell replacement λ. We assume that methylation occurs independently across each allele, so that the transition rate from the double-methylated to the single-methylated state is twice that from the single-methylated to the demethylated state (and similarly for the demethylation rate). However, in future PHYFUM versions, we will relax this assumption by introducing two additional parameters. We use standard matrix exponentiation to calculate the transition probability matrix along the branches, given the initial states, the three rates described above, and a specified time interval. This likelihood function is directly used to calculate the phylogenetic likelihood using a minor modification of Felsenstein’s pruning algorithm^8^ for somatic evolution (described below). Finally, the total likelihood is calculated by multiplying the likelihoods of all fCpGs, assumed to evolve independently.

#### 1.2. Error model

The observed data (continuous methylation fractions *β*) need to be mapped to PHYFUM’s discrete substitution model states (each with its expected methylation fraction 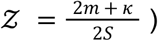. We use the likelihood of each model state under a simplified version of the error model in the original *Flip-Flop* model implementation^2^. Thus, each *β* is a mixture of 𝒵 beta-distributed random variables with means equal to a linear transformation of 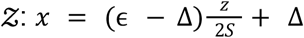 that accounts for background noises that offset the minimum Δ and maximum ϵ observable *β*, and a shared scale parameter *κ* that controls the breadth of the distributions and thus models the assay’s precision. The three error-model parameters are inferred by PHYFUM.

#### 1.3. Tree model

The tree model describes evolution at the stem-cell-niche level. Thus, internal nodes correspond to stem-cell-niche division events, and the tree shape reflects stem-cell-niche (gland) demography. It differs from the standard tree model in some aspects of the phylogenetic likelihood calculation and in the different gland-division models.

##### 1.3.1. Somatic phylogenetic likelihood calculation

Under the default gland-division model, the phylogenetic likelihood is calculated by integrating over all possible states at internal nodes following the standard Felsenstein pruning algorithm^8^. The tree is traversed in postorder (starting at the tips towards the root), calculating the partial likelihoods *L*_*A*_ (*a*), or probability of observing state *a* at node *A* given its descendant nodes *X* and *Y* with branch lengths *t*_*X*_ and *t*_*Y*_ and Ω the set of 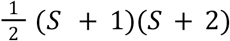 states in the fluctuating methylation model:

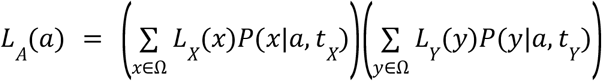

At the tips, the partial likelihoods *L* are given by the error model and otherwise calculated recursively using this equation, and the transition probabilities *P* are calculated using the fluctuating methylation model. When the likelihood calculation reaches the MRCA traversing the tree in post-order, the LCA partial likelihoods are calculated equivalently for a 1-degree node:

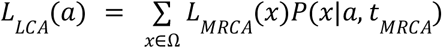

This additional LCA node allows us to use non-reversible CTMC models and incorporate biological knowledge of the origin of the somatic evolutionary process, as we did previously in PISCA^7^.The overall likelihood of a single fCpG is terminated by summing over the products of the partial likelihoods and their prior state probability at the LCA π:

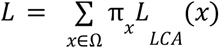

By default, π applies half of the weight to the fully methylated and fully demethylated states, reflecting the elimination of methylation patterns inherited from paternal gametes in the early stages of embryogenesis. This is a departure from the original pruning algorithm, in which the CTMC is reversible and assumed to be in equilibrium, π denotes the equilibrium frequencies, and this termination can happen anywhere in the tree.

##### 1.3.2. Gland-division models

The default model, *Cloning*, in which dividing glands generate exact clones, is computationally convenient but could be biologically unrealistic. We devised three alternative models: *Budding, Double-split*, and *Split-double*, which allow us to test how glands divide using Bayesian model comparison. In all cases, we assume that the stem-cell niches recover their fixed size instantaneously after division. We always integrate over all possibilities of stem-cell and daughter-lineage assignment with equal probability and implement the different models by modifying the partial-likelihood calculation only. For estimation (but not for simulation unless noted), we also make the simplifying assumption that these divisions occur only at internal nodes, not along the branches.

First, we simplify the notation of the partial likelihood calculation equation by defining the daughter-conditioned partials φ _*c*_ (*a*).

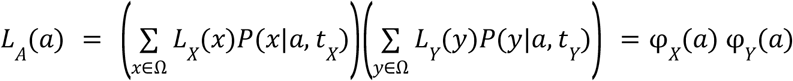

Then, we generalize it to consider a set of possible combinations of daughter ancestral states *C*(*a*) ⊆ Ω *x* Ω, and their probability *P*(*b, c*|*a*) of being generated from the ancestor state *a*. We also explicitly treat states as vectors indicating the unique combination of the numbers of unmethylated, half-methylated, and fully methylated cells they correspond to ***a*** = (*a, a, a*,) = (*S* − *k* − *m, k, m*).

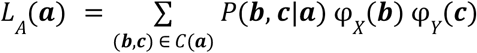

Biologically, this represents a gland division in which the resulting offspring states may differ from the ancestral state. In the *Cloning* model, this is not allowed, and thus *C*(***a***) = {(***a, a***)} and *P*(***b, c***|***a***) = 1, defined only on *C*_*model*_(***a***), collapsing to the standard partial likelihood calculation. The different stem-cell niche division models generate different sets of combinations of allowed ordered daughter states *C* (***a***) and their probabilities *P*_*model*_ (***b, c***|***a***) defined only on that support.

*Budding* models a single stem cell generating a daughter gland by producing *S* identical copies of itself, while the original gland remains unchanged. Therefore, the set of allowed ordered daughter-state combinations *C* (***a***) consists of pairs with a daughter in the ancestral state ***a*** and the other in a ***d***_*r*_ state with all *S* cells assigned to a single methylation state category *r* present in ***a***:

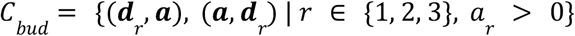

The probability of this *r* category is proportional to its count in the ancestral state 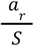, and each daughter lineage has a probability of undergoing budding that is typically equal between lineages (*Pl*_*X*_ = *Pl*_*Y*_ = 0. 5):

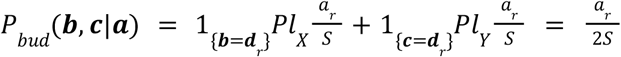

In PHYFUM, this is implemented as a pre-computed Ω *x* 3 matrix of budding probabilities, pre-computed states for buds originating from each of the 3 possible stem-cell types, and a partial-likelihood calculation that sums partials for the three possible buds in each of the two lineages, and then calculates their mixture.

*Double-split* models a perfect duplication of the stem cell niche (i.e., every stem cell makes an identical copy of itself), followed by their distribution in two *S* stem-cell niches at random. The probability of obtaining a given state ***b*** = (*b*_1_, *b*_2_, *b*_3_) given the ancestral state ***a*** for a given daughter niche is given by the multivariate-hypergeometric mass:

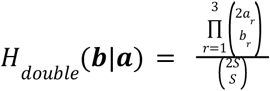

The set of allowed ordered daughter-state combinations *C*_*double*_ (***a***) consists of all combinations of daughter states for which the sum of their cells in each category is twice that of their ancestor niche.

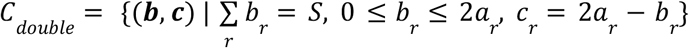

To calculate the probability of a combination of allowed daughter-state combinations, we assume equal probability of each daughter being sampled by the multivariate-hypergeometric distribution (*Pl*_*X*_ = *Pl*_*Y*_ = 0. 5):

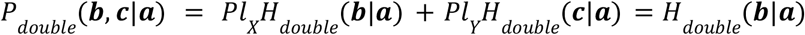

In PHYFUM, we implemented this model using a pre-computed Ω *x n* (*C*_*double*_) matrix of gland-division probabilities and a Ω *x n*({*X, Y*}) *x n* (*C*_*double*_) matrix of states in each daughter and gland-division combination, which are used to compute the *L*_*A*_ (*a*) sum over (*C*_*double*_).

*Split-double* models the division of the ancestral stem cell niche into two niches of ⌈*S*/2⌉ and ⌊*S*/2⌋ cells. The larger niche ***u*** = (*u*_1_, *u*_2_, *u*_3_) is seeded by a multivariate hypergeometric sampling of ***a***, with probability mass function:

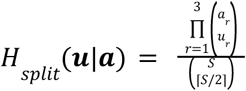

After which ***u*** and its complementary niche duplicate. For even *S*, this restores both daughter niches to size *S* and their states to the final ***b*** = 2***u***, and ***c*** = 2(***a*** − ***u***). When *S* is odd, the round of duplication after the split produces niches of size *S*+1 and *S*-1, which we assume are followed by the death or duplication (respectively) of a single cell (with probability proportional to its category counts) to restore both daughters to size *S*. The resulting set of ordered daughter states *C*_*split*_ (***a***) that arise from a composition of the multivariate hypergeometric sampling, duplication, and stochastic single-cell death and duplication is:

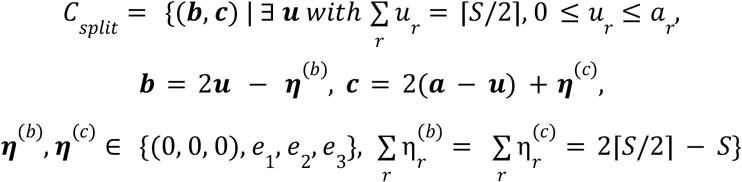

With *e*_1_ denoting the unit vector in category *r*. For each (***b, c***) ∈ *C*_*split*_ (***a***) there exists a unique intermediate state ***u*** = *u*(***b, c***). Finally, the split probabilities are defined by:

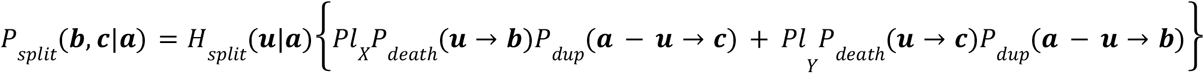

Where the probabilities of the adjustments to size *S* via cell duplication and death are:

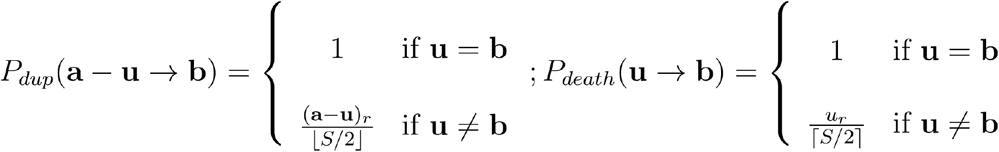

With *r* being the cell category that undergoes the adjustment event (i.e., *r* : *u*_*r*_ ≠ *b*_*r*_).

In PHYFUM, the *Split-double* implementation follows the same strategy as the *Double-split* one, but for *C*_*split*_ instead.

#### 1.4. Bayesian model comparison and averaging

##### 1.4.1. Stem-cell-niche size

The number of stem cells in the niche *S* determines the dimensions of the rate matrix and the number of expected methylation rate peaks in the error model. Allowing this parameter to vary within the MCMC changes the dimensionality of the substitution model, complicating its implementation and likely leading to mixing issues. Instead, PHYFUM is run separately for a given set of fixed *S*, and the best-fit *S* is found via Bayesian model comparison, or the conditional posteriors are marginalised using Bayesian model averaging. PHYFUMflow automates both processes.

Bayesian model comparison is typically computationally expensive, as it requires sampling from a series of power posteriors to estimate marginal likelihoods using stepping stone^14^ or path sampling^15^. Alternatively, the harmonic mean estimator only requires samples from the posterior, but it can fail catastrophically when its variance becomes very large. For the specific purpose of selecting PHYFUM’s optimal *S*, these strategies perform similarly in both biological and *in silico* data (Supplementary Note 1). All of them are supported by our suite of programs.

Instead, if the marginalised posteriors are the objective, we marginalise *S* via model averaging by resampling the conditional posteriors with replacement. For data *D*, continuous parameters *𝞡*, and the discrete parameter *S*, the joint posterior is:

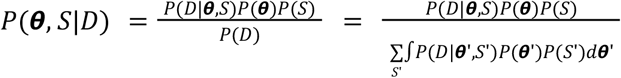

From PHYFUM, we obtain conditional posteriors on fixed *S, P*(*𝞡*|*D, S*) and their conditional marginal likelihoods *P*(*D*|*S*):

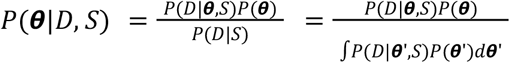

We calculate the posterior probability of each value of *S* by marginalising the joint posterior over *𝞡*:

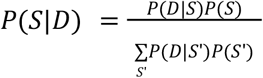

Finally, we average over *S* to obtain the marginalised posterior of interest:

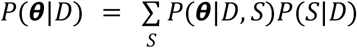

This identity follows directly from 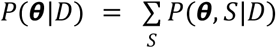 and is standard in Bayesian phylogenetic model averaging.

In practice, we use a uniform prior *P*(*S*) and estimate 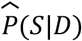 using a small subset 𝓈 of all possible *S* values 𝒮, 𝓈 ⊂ 𝒮, *s* ∈ 𝓈, ensuring that 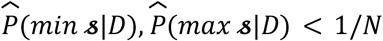 with *N* being the number of posterior samples desired in the marginalised sample, and that the function *s* ⟼ *P* (*D* | *s*) for *s* ∈ 𝓈 is unimodal and monotonic in each tail. Otherwise, we expand 𝓈 as needed.

Importantly, *S* does not alter the parameter space and thus could theoretically be estimated using BEAST’s standard Metropolis-Hastings MCMC algorithm in future versions of PHYFUM.

##### 1.4.2. Gland-division model comparison

Currently, PHYFUM can only use a single gland-division model per MCMC run, as for *S* above. Therefore, we resort to Bayesian model comparison to select the best-fit gland-division model and to calculate its goodness of fit. PHYFUM’s marginal likelihood estimation (using the harmonic mean estimator, stepping stone^14^, or path sampling^15^) yields estimates of the marginal likelihood conditional on a combination of stem-cell-niche size *S* and gland-division model *M P*(*D*|*M, S*).

Typically, we are interested in the model-fit marginalised over *S*. For which, for each model, we first calculate the model evidence *P*(*D*|*M*) by integrating *S* out of the conditional marginal likelihoods with its prior:

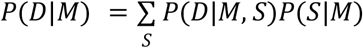

In practice, for each model, we use the same uniform prior *P*(*S*|*M*) and integrate it over a small subset 𝓈 of all possible *S* values 𝒮, 𝓈 ⊂ 𝒮, *s* ∈ 𝓈 similarly to the model averaging over *S* above. Here, we ensure that ensure 𝓈 is the same for all models, expanding it if needed, until *BF*_*bestfit* ***s***, *min* ***s***_, *BF*_*bestfit* ***s***, *max* ***s***_ ≥ 10 for all models to ensure the omitted contribution to 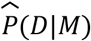 is small.

With the model evidence 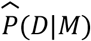 estimated, we perform standard Bayesian model comparison using Bayes factors and calculate model posteriors by marginalising the joint posterior over *𝞡*:

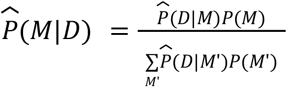

Typically, we assume a uniform prior among the models considered.

Of note, for the specific purpose of selecting the gland-division model, the harmonic mean estimator performs very similarly to the more computationally expensive stepping stone^14^ or path sampling^15^ methods *in silico* (Supplementary Figures 15-16). Our suite of programs supports all of them.

## 2. Simulation studies

To evaluate PHYFUM’s accuracy under different conditions, we developed a phylogenetic fCpG simulator under the PHYFUM model. Given all parameter values (including the fluctuating methylation model, prior state probability at the LCA, noise model, and gland-division model) and a chronogram (or parameter values to generate it under the coalescent model), this simulator evolves the specified number of fCpG sites independently along the tree to then generate their final methylation fractions. The *Budding* gland-division model differs slightly between simulation and estimation. The estimation model assumes the bud originates by cell division, and thus the ancestral stem-cell-niche remains unchanged, while the simulation model assumes the bud originates by cell migration, and thus a single cell replacement happens in the ancestral stem-cell-niche. Importantly, in simulation, stem-cell-niche divisions can also be simulated along branches at a user-specified rate, whereas the estimation model considers them only at internal nodes.

### 2.1 Broad full-factorial simulation study

We explored PHYFUM’s behaviour on a very broad parameter space. We generated 486 simulations with 2,000 fluctuating CpG sites and all combinations of several parameter values:

- Stem cells, *S*: [3, 10, 20]
- Stem cell replacement rates, λ: [0.1, 1, 10]
- Methylation rate, *μ*: [0.001, 0.01, 0.1]
- Demethylation rate, *γ*: [0.001, 0.01, 0.1]
- Assay precision, *κ*: [20, 50, 100]
- Phylogenetic tree: [fixed 4-sample tree, fixed 9-sample tree]
- Sample age = 45
- Maximum observable methylation fraction *β*, ϵ = 0.93
- Minimum observable methylation fraction *β*, Δ = 0.05
- Gland-division model: *Cloning*

We ran three independent 750k-iteration MCMC estimation chains per condition, sampling every 75, and discarded the first 10% of each as burn-in, with all other PHYFUM parameters as default in PHYFUMflow unless specified. We ran the estimation for each condition with stem niche size (estimation S, *eS*) 3 and 10. We did not run *eS* = 20 due to the super-quadratic computation time scaling with *eS*.

For the data analysis, we considered an estimation valid (i.e., properly mixed and converged) if the effective sample sizes (ESSs) ≥ 200 and the Potential Scale Reduction Factor (R-hat) ≤ 1.05 for all continuous parameters across the three chains and their combination, independently of the final number of samples in each (if interrupted due to running-time limits). For the tree, we calculated the normalised RF (Robinson-Foulds) and normalised wRF (weighted Robinson-Foulds) distances^9,16^ between each sampled tree and the true tree, and the ESS and R-hat of those continuous variables, following the pseudoESS strategy^17^. We calculated the total estimation error per estimation condition as the sum of the weighted RF distance and the relative errors of the methylation, demethylation, and cell-replacement rates.

We estimated the effect of the different simulation parameters on accuracy, restricted to estimations with *eS = S*, using random forests’ percent increase in mean squared error and linear model’s coefficients after scaling the predictors. For this, we considered all variable simulation parameters and the following derived statistics: evolutionary rate (GM(*λ,μ,γ*), GM = geometric mean), methylation imbalance (*μ*/γ), and cell-replacement-methylation imbalance (*μ*/*λ*). For the predictors with the strongest effect, we compared the distribution of total error across predictor levels using the Kruskal-Wallis rank-sum test followed by the post hoc Dunn test, and report p-values adjusted for multiple comparisons using the Holm method. To study the effect of *eS* in the estimation of each parameter, we calculated the difference in relative errors between *eS* = 3 and *eS* = 10 for each simulation condition without imbalanced methylation, and assessed if these differences were distributed around 0 using a one-sample Wilcoxon signed-rank test per *S* with Holm correction for multiple testing (for the multiple *S* within a parameter). Additionally, we removed conditions with (de)methylation rates > 0.1 to control for phylogenetic saturation.

### 2.2 Targeted simulation study

To characterize PHYFUM’s accuracy under realistic evolutionary parameters, we conducted a simulation study in which the evolutionary rate varies across a wide range while methylation and demethylation are highly correlated, as observed in real data after fCpG selection, and the relationships of both with the cell-replacement rate are also controlled. We fixed other parameters that had minor effects on the accuracy in the previous simulation study, and considered the different gland-division models. We generated 600 simulations out of 50 conditions simulated along 150 coalescent trees per gland-division model with 2,000 fluctuating CpG sites each:

- Stem cell replacement rates, λ: [0.5, 0.75, 1.0, 1.25, 1.5]
- Methylation rate, μ: λ *x*_*μ*_ ^−1^, *with x*_*μ*_ *taking* 10 *evenly spaced values between* 10 *and* 500
- Demethylation rate, γ: λ *x*_*γ*_ ^−1^, *with x*_*γ*_ *taking* 10 *evenly spaced values between* 8 *and* 450
- Phylogenetic tree: coalescent 5-sample tree (sampled per replicate)
- Gland-division model: [*Cloning, Double-split, Split-double, Budding*]
- Stem cells, *S* = 6
- Sample age = 45
- Assay precision, *κ* = 100
- Maximum observable methylation fraction *β*, ϵ = 0.92
- Minimum observable methylation fraction *β*, Δ = 0.04

We estimated each condition across the four gland-division models using the true *S* (2,400 estimation conditions), following the same running and QC protocol as in the broad full-factorial simulation study, unless otherwise specified. For most estimation conditions, we ran two independent MCMC chains, followed by marginal-likelihood estimation for each (sampling 100 power posteriors with the same total number of iterations as the MCMC, with powers ∼ beta(0.3,1)). For gland-division model combinations with a higher rate of QC issues, we ran a third chain and discarded the chain with the smallest posterior only if QC passed with the other two chains. Otherwise, all three were kept, and the estimation was considered invalid. We calculated the posterior probability of each model per condition, estimation, and simulation model, as described in 1.4.2, using stepping stone marginal likelihoods for the main analyses and the harmonic mean estimator for comparison.

To compare PHYFUM against our previously published single-cell fCpG model implemented in PISCA, we estimated the *Cloning* simulations in PISCA using three independent 50M MCMC chains per condition, sampling every 10k, and discarded the first 10% of each as burn-in. We used PISCA’s default parameters, with settings equivalent to PHYFUM’s unless specified here. The constant effective population size was initialized at 10. The heterozygous methylation rate was initialized at μ, with a prior ∼ N(0, 10*μ). The other three rate parameters (heterozygous demethylation, homozygous methylation, and homozygous demethylation) relative to the heterozygous methylation rate were initialized at 1 with a prior ∼ logN(1,0.6). We followed the same running and QC methodology as in the broad full-factorial simulation study. Since the rate parameters are not directly comparable with PHYFUM, we compared their accuracy under conditions with valid estimates for both methods using a paired Wilcoxon signed-rank test on their tree reconstruction errors (wRF).

We characterized PHYFUM’s gland-division model classification performance using multiclass one-vs-all ROC curves and a confusion matrix. In the multiclass one-vs-all ROC, each model is used as the reference in turn. We use the data for all estimations under the reference model (simulated under any model), with their posterior probability as the classification score, and the match between the simulation and estimation models as the positive class. The confusion matrix shows the proportion of conditions simulated under a model for which a given model is the best fit. We considered only simulation conditions with valid estimates for all combinations of estimation and simulation models. Removing this QC filter does not qualitatively change the results.

We characterized the relationship between model distinguishability and conditional accuracy by calculating the accuracy after discarding conditions for which the second-best-fit model was closer than a sweeping Bayes Factor threshold to the best-fit model (from 0 to the maximum observed), and plotted the conditional accuracy against the proportion of conditions discarded. We also tested if model distinguishability decreases with the evolutionary distance to the last modeled gland division by testing if there was a rank correlation between the probability of *Budding* and the evolutionary distance from the most recent internal node to the present (*t*λ + *tμ* + *tγ, t* = time to the present) for all simulations in which *Budding* was not the true model and its estimation passed QC (i.e., we tested if there was an increase in the probability of the wrong model with the increase in evolutionary distance to the last modeled event).

## 3. Phylogenetic reconstruction on patient glands

### 3.1. DNA methylation profiling

Methylation fractions were obtained per our previous work^2^. Initial analysis of our previously-derived fCpGs revealed that these sets underwent epigenetic switching at too high a rate to properly calibrate the phylogenetic clock rate, showing signs of phylogenetic saturation, so we derived new sets without these issues. Inspired by our recent work^4^, we employed the Laplacian score to remove tissue-specific signal and CpGs which systematically varied between patients. To do this, we repeatedly (n=100) selected a single sample per patient and calculated the laplacian score for each CpG and the mean intra-tissue CpG standard deviation. We retained CpGs which had:

1. High mean intra-tissue standard deviation (>0.15), indicating high heterogeneity across the cohort.
2. High mean Laplacian score (>0.8), ensuring minimal tissue or patient-specific signal.
3. Average mean methylation between 0.4 and 0.6, ensuring approximately equal forward and backwards epigenetic switching rates.

The remaining 16,219 CpGs were considered as fluctuating and were used for inference (Supplementary Figure 17).

### 3.2. Single-sample evolutionary inference

For comparison against PHYFUM, unless specified otherwise, we used the original *Flip-Flop* model from our previous work^2^ to estimate evolutionary parameters using the new sets of fCpG sites. The original *Flip-Flop* model differs from PHYFUM’s in its error model, whereby each beta-peak was assigned its own precision parameter, κ_*z*_, rather than having a global precision parameter *κ*. To show the effect of changing the error model on the estimation of *S*, we also used a new version of the *Flip-Flop* model that shares PHYFUM’s error model, although it is not strictly equivalent since it models heterozygous and homozygous (de)methylation separately, as in our recent work^4^.

### 3.3. Phylogenetic inference

Following fCpG selection, we used PHYFUMflow to generate an input file for PHYFUM per patient, tissue, *S*, and gland-division model, using their fCpG and age data. We ran two independent 2.5M-iteration MCMC estimation chains for each combination of *S* and gland-division model, sampling every 250, and discarded the first 10% of each as burn-in, with all other PHYFUM parameters set to default in PHYFUMflow unless specified. Initially, we explored *S* = [4,10] for all tissues except the endometrium, for which we used a larger initial range, *S* = [4,15], based on previous results. We expanded these estimation intervals as needed up to *S* = 20 to marginalize over *S* (see Methods 1.4.1) and compared gland-division models using harmonic mean estimates of the marginal likelihood (see Methods 1.4.2). When S > 15, we used 1M-iteration MCMC chains sampled every 100 except for the EAN_Colon patient, for which we improved mixing by increasing the weights on the operators which modify Δ, ϵ, *β*, and λ, and ran 0.5M-iteration MCMC chains sampled every 50. A single patient (EAN_Colon) was eliminated from the statistical analyses across the gland-division models since its results could not be marginalized over *S* reliably within the estimation range for some of the alternative models. For the data analysis, we considered an estimation valid (i.e., properly mixed and converged) if the effective sample sizes (ESSs) ≥ 200 and the Potential Scale Reduction Factor (R-hat) ≤ 1.05 for all continuous parameters in the combination of the estimation chains, independently of the final number of samples in each (if interrupted due to running-time limits).

Flatter-than-usual posteriors are associated with weak phylogenetic signal and may indicate estimation issues. We checked whether this issue was present by comparing the density defining the 0.95 highest-density region in the posterior with the maximum density in the prior for the three stem-cell-niche rate parameters, and deemed it present if the second was larger than the first for all three rates. We used the posterior samples of the best-fit *S* per patient for this test. Only patient UQ was discarded for this reason for all biological data results related to Figure 2. This issue is also reflected in unusually wide HDIs (Supplementary Figure 8). These results do not change qualitatively contingent on this QC step, except for the comparison of rates between tissues.

Inference on most patient and simulated data was carried out on the Sol supercomputer^18^, administered by Research Computing at Arizona State University.

### 3.4. Statistical analyses

To compare PHYFUM’s results across tissues, we tailored the statistical tests to the specific characteristics of each parameter. We used a Dunn test to compare relative MRCA ages. For each stem-cell-niche parameter and absolute MRCA age, we used linear regression models including the reciprocal of age as a covariate (except for *S*, for which we did not inverse age). We used the *emmeans* R package to query the marginal means per tissue. We corrected for multiple tests per parameter using the Holm correction in all cases.

We compared PHYFUM and *Flip-Flop* estimates using Pearson’s correlations between the estimates (mean across patients and tissues for *Flip-Flop*). We also performed this comparison using a different version of *Flip-Flop* that uses the same error model as PHYFUM but differs in the stem-cell-niche model formulation. To investigate the observed weakened associations between PHYFUM and *Flip-Flop* parameter estimates in the endometrium, we used linear regression models of λ on *μ* + *γ* per tissue and estimation model.

We compared gland division modes by calculating model posteriors per patient and tissue (see Methods 1.4.2) and Bayes factors against the best-fit model. We refer to a Bayes factor >10 as strong and >100 as decisive.

