## Supplementary Materials for "PHYFUM: Phylogenetic Reconstruction of Normal and Pre-malignant Tissue Evolution Using Fluctuating Methylation"

Supplementary Figures ... 2
Supplementary Tables ... 15
Supplementary Methods ... 16
1. PHYFUM: additional implementation details ... 16
2. PHYFUMr ... 16
3. PHYFUMflow ... 17
4. PHYFUM simulator ... 18
Supplementary Note 1 ... 19
Effect of marginal likelihood estimation method in Bayesian model comparison for stem-cell number estimation ... 19

### Supplementary Figures

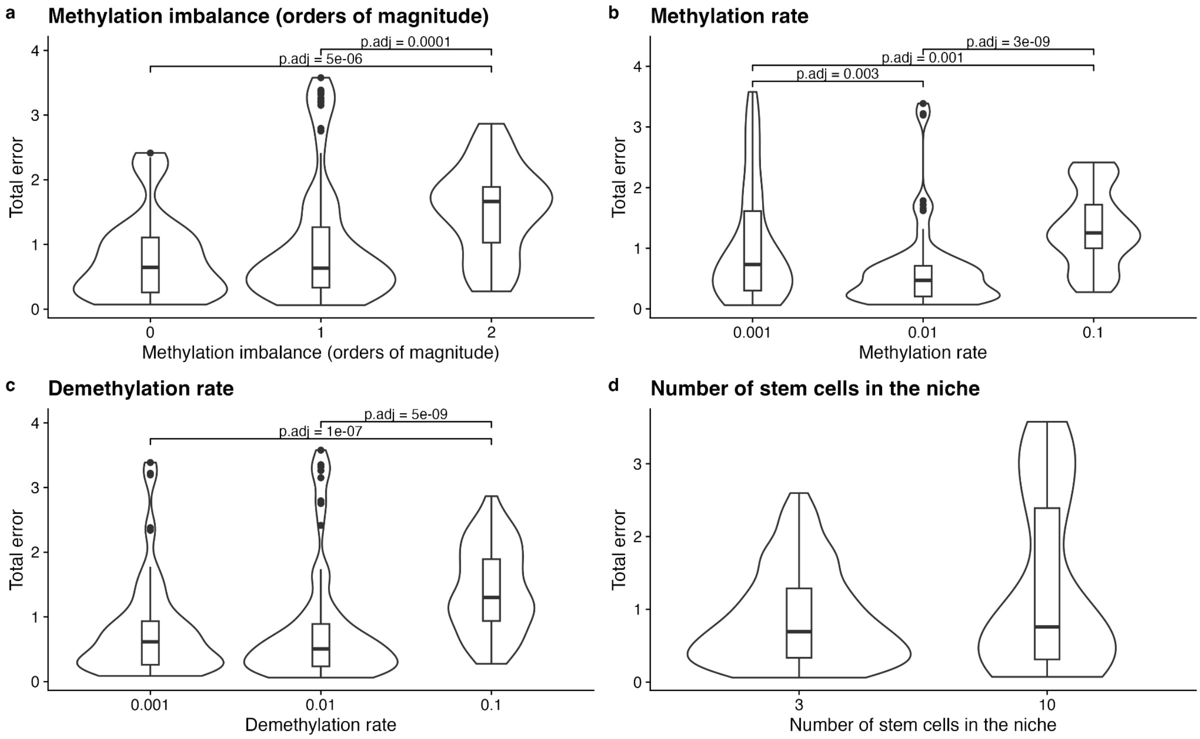


**Supplementary Figure 1: Impact of influential simulation parameters on estimation accuracy.** Distribution of the sum of relative errors of methylation, demethylation, and cell-replacement rates, and weighted RF distance in estimates of the broad full-factorial simulation study under the true stem cell niche size. Dunn test with Holm-adjusted p-values for significant contrasts.

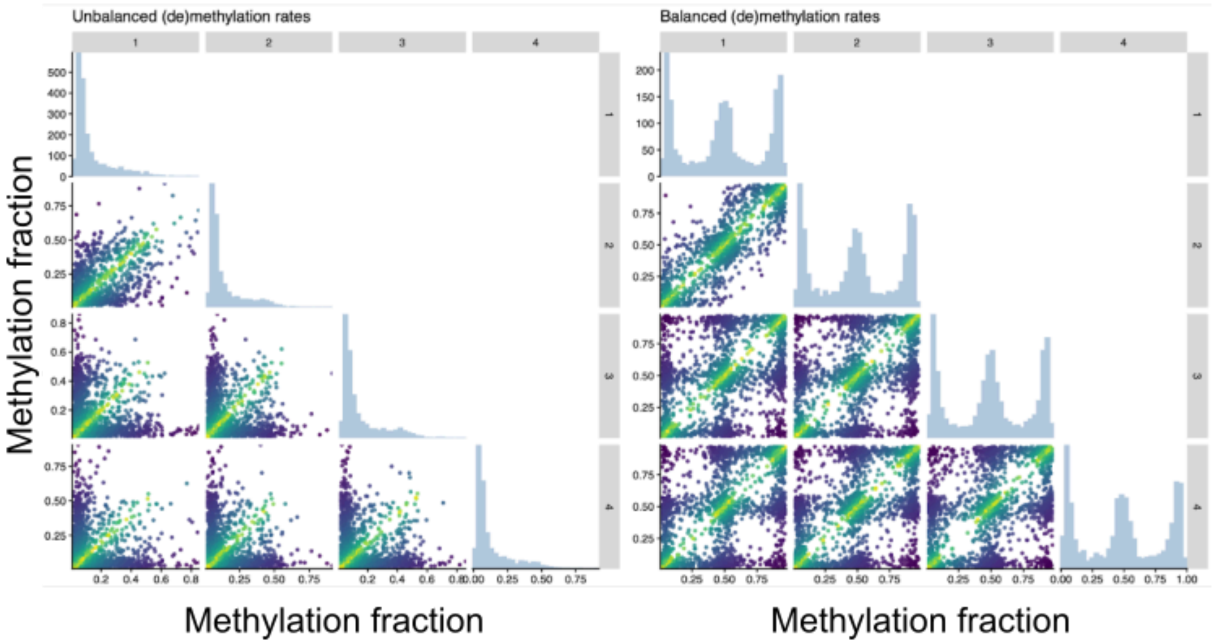


**Supplementary Figure 2: Imbalanced methylation generates clear patterns in the distribution of methylation fractions.** Representative example of a simulated patient with four samples undergoing imbalanced and balanced epigenetic modification rates. Methylation and demethylation rates differed by one order of magnitude in the first case (0.01 and 0.1 mutations per year, respectively), and were equal at 0.01 mutations per year in the second. The remaining parameters, including tree topology, are shared across the simulations and lie within the optimal parameter space.

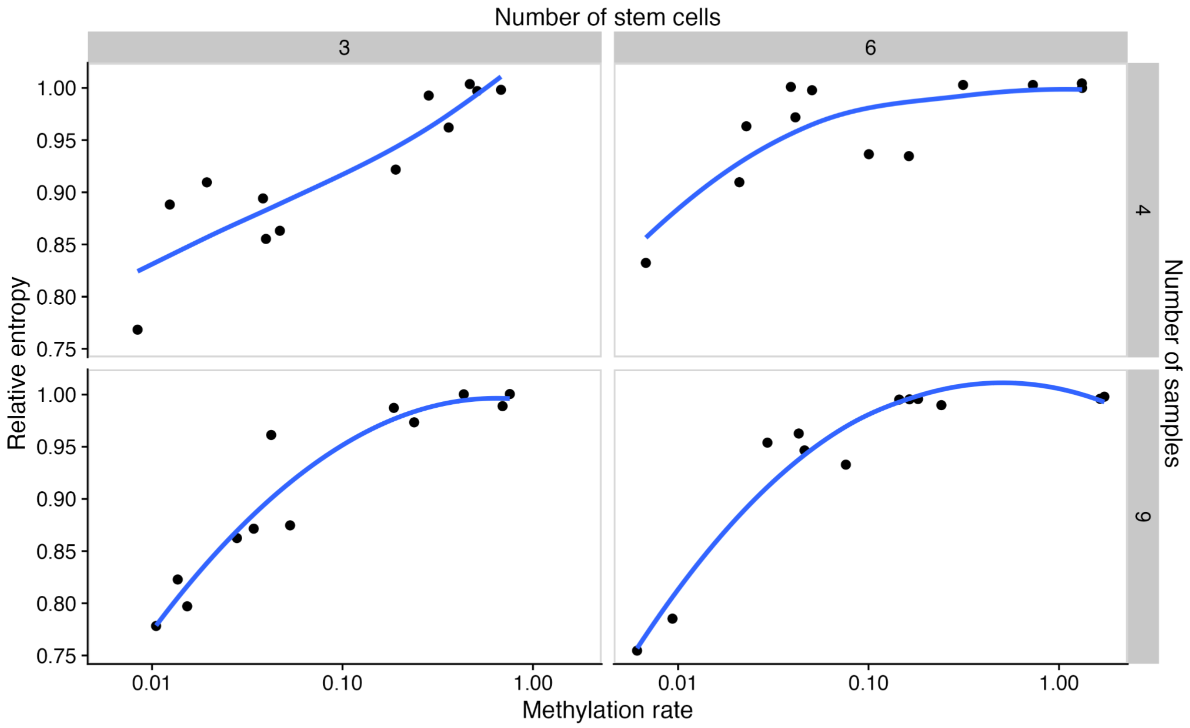


**Supplementary Figure 3: Saturation detection using relative entropy.** Relationship between the relative entropy statistic and the increase in switching rate that drives saturation. Local regression curve in blue.

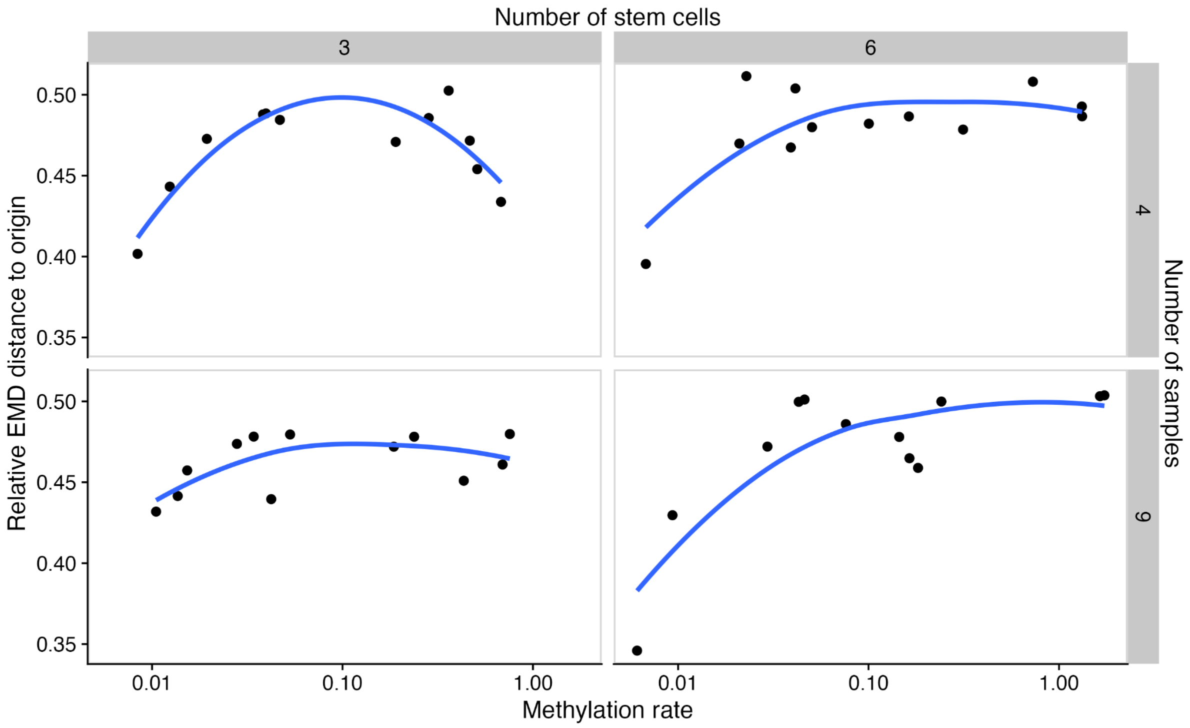


**Supplementary Figure 4: Saturation detection using the relative earth moving distance of frequency distributions.** Relationship between the relative earth-moving distance between the observed empirical frequencies and the LCA, and the increase in switching rate that drives saturation. Local regression curve in blue.

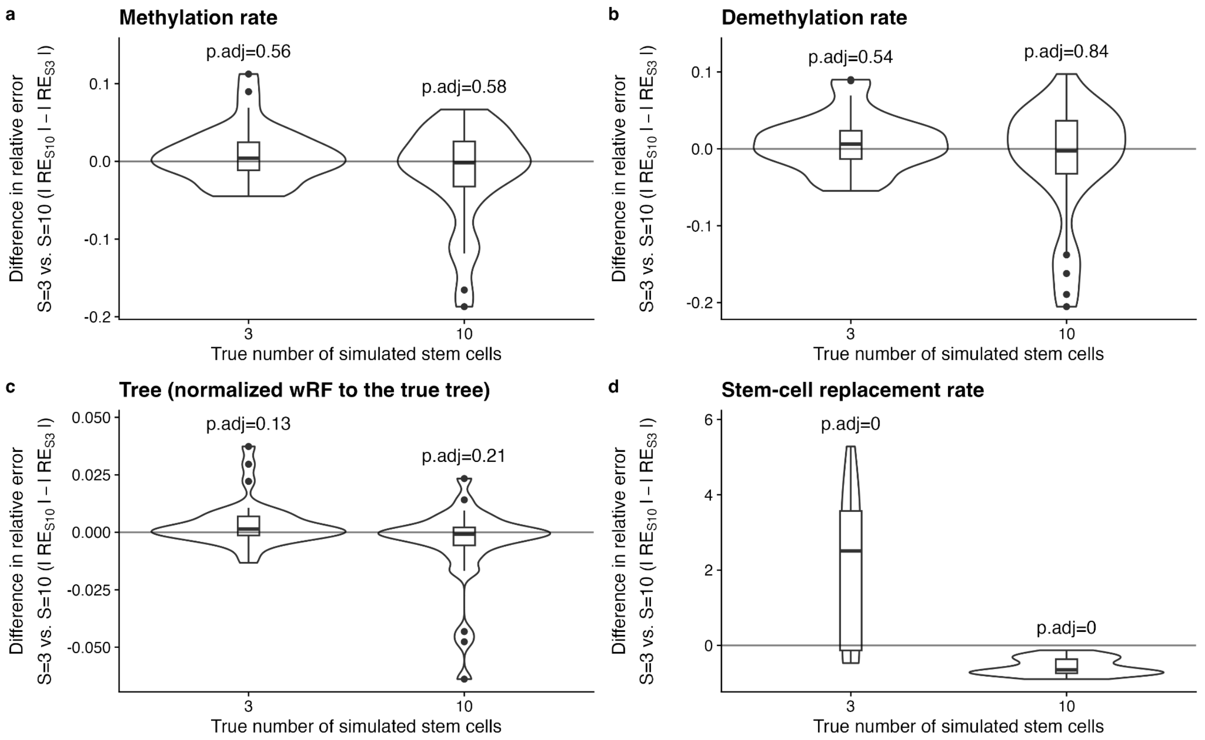


**Supplementary Figure 5: Negligible effect on PHYFUM's estimation accuracy when assuming a wrong stem-cell-niche size without imbalanced methylation and saturation.** Distribution of the difference in absolute relative errors between estimates carried with size 10-stem-cell niches and 3-stem-cell niches for data simulated under 3 and 10-cell niches. Positive numbers indicate that 3-cell estimates were more accurate. Methylation (a), demethylation (b), and trees (c) are better estimated, on average, under the true stem-cell-niche size. Still, the difference in accuracy is small and not statistically significant from 0 (one-sample Wilcoxon signed-rank test with Holm correction for multiple testing within parameters). In contrast, the stem-cell replacement rate (d) scales with the number of stem cells in the niche (a higher number of cells requires a higher replacement rate to reach fixation in the same time) and thus is strongly influenced by this parameter. Broad full-factorial simulation study conditions with methylation rate = de-methylation rate < 0.1, to avoid the two largest easily avoidable confounding sources of errors (imbalanced methylation and saturation).

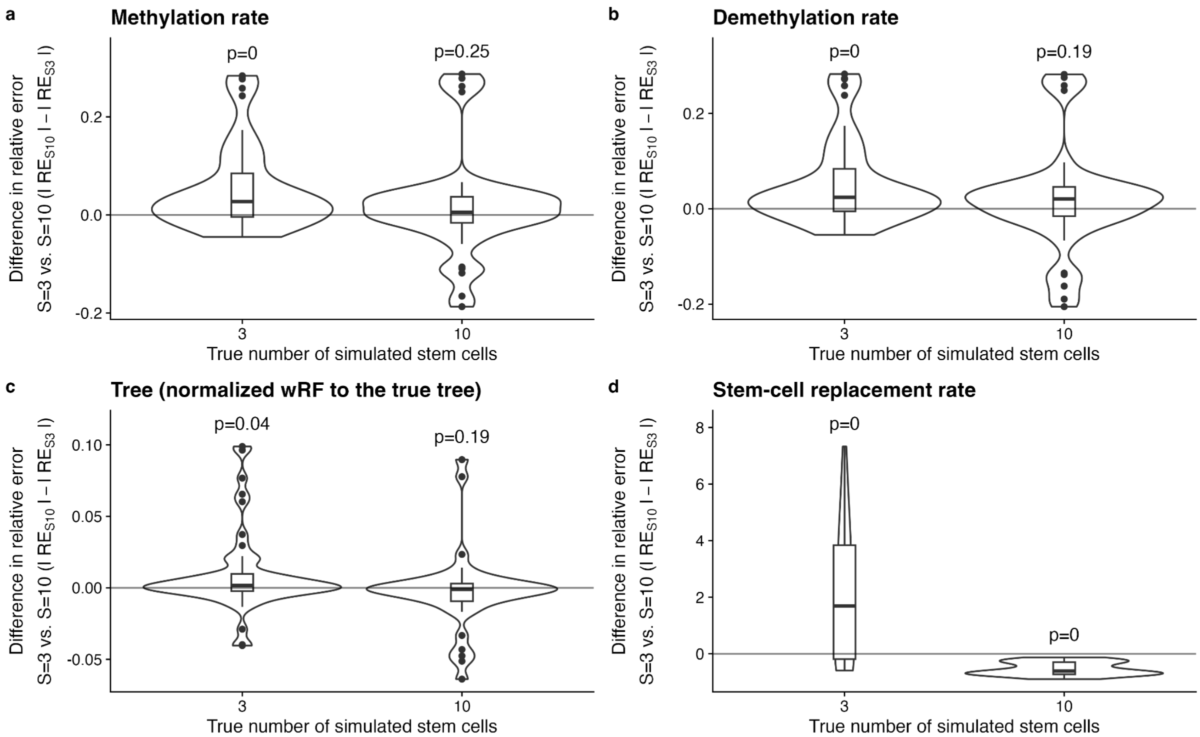


**Supplementary Figure 6: Negligible effect on PHYFUM's estimation accuracy when assuming a simpler model (smaller stem-cell-niche size) without imbalanced methylation.** Distribution of the difference in absolute relative errors between estimates carried with size 10-stem-cell niches and 3-stem-cell niches for data simulated under 3 and 10-cell niches. Positive numbers indicate that 3-cell estimates were more accurate. Methylation (a) and demethylation (b) are better estimated, on average, under the smallest stem-cell-niche size, but this difference is only statistically significant when the smallest stem-cell-niche size is the truth (one-sample Wilcoxon signed-rank test with Holm correction for multiple testing within parameter). Tree reconstruction is more accurate under the true stem-cell-niche size, but the difference is only statistically significant for the true small *S*. In contrast, the stem-cell replacement rate (d) scales with the number of stem cells in the niche (a higher number of cells requires a higher replacement rate to reach fixation in the same time) and thus is strongly influenced by this parameter. Broad full-factorial simulation study conditions with methylation rate = de-methylation rate, to avoid a confounding source of error not present in biological data (imbalanced methylation).

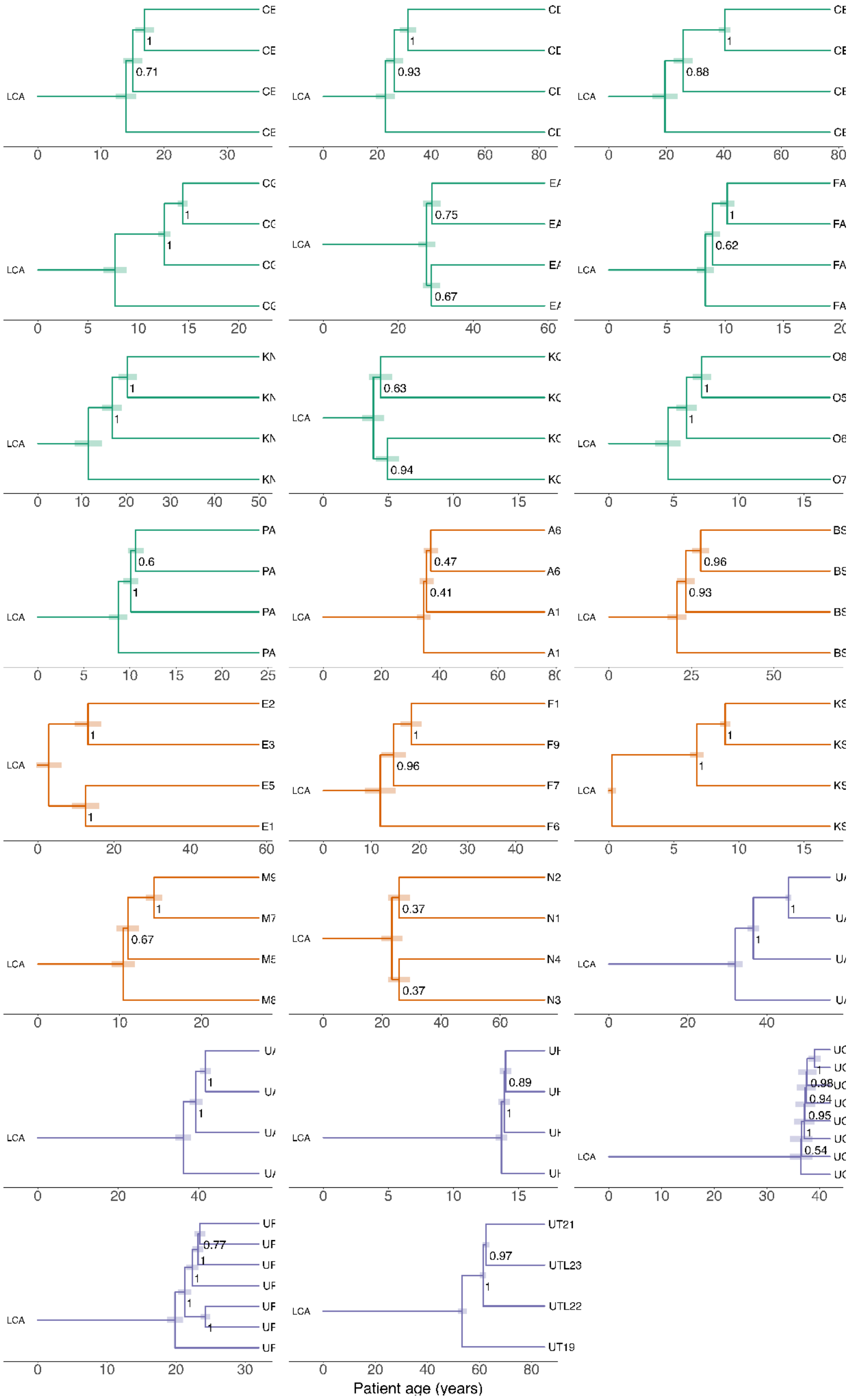


**Supplementary Figure 7**: **Phylogenetic trees of biological samples.** Inferred maximum clade credibility (MCC) trees on biological samples of colon (green), small intestine (orange), and endometrium (purple). The shaded boxes spanning the nodes highlight the 95% HPD. Node labels show posterior probabilities.

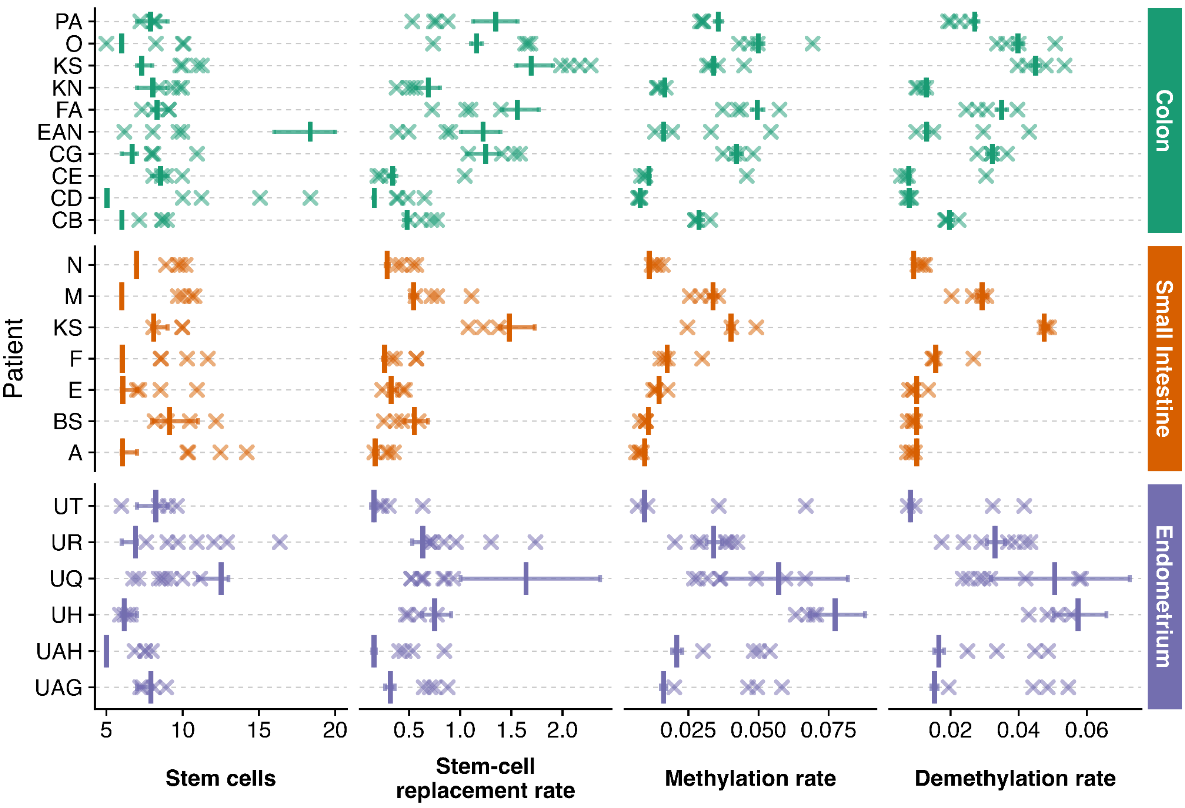


**Supplementary Figure 8**. **Comparison between parameter estimates with PHYFUM and the *Flip-Flop* method**. Single-sample estimates are represented with crosses; 95% High Density Intervals (HDI) are depicted with coloured horizontal lines, and the posterior means are shown with vertical coloured bars.

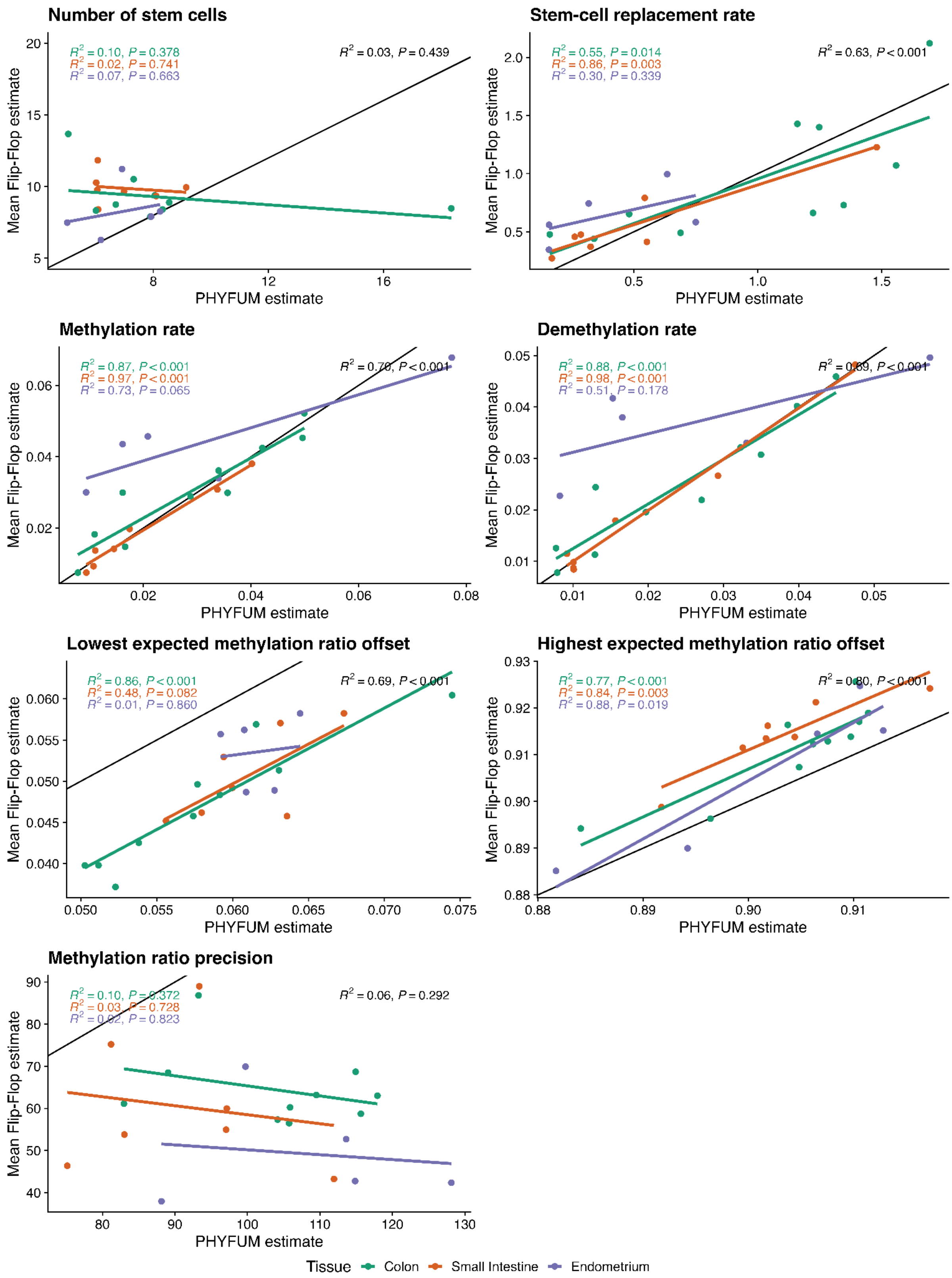


**Supplementary Figure 9: Linear relationships between parameter estimates (stem-cell niche and error models) using the original *Flip-Flop* model implementation and PHYFUM**. Note that the methylation ratio precision (scale parameter of the beta distribution) is not equivalent between the two error models since *Flip-Flop* uses a different precision per methylation ratio expected peak (of which here we report the mean) while PHYFUM uses a single scale for all beta distributions.

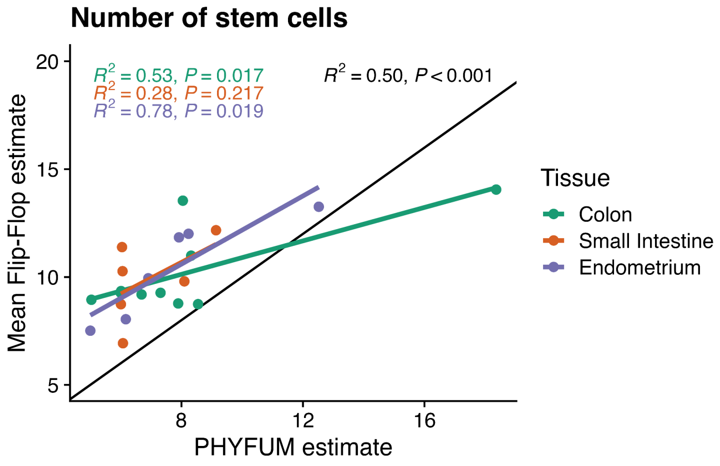


**Supplementary Figure 10: Linear relationship between the estimated number of stem cells between PHYFUM and an alternative implementation of the *Flip-Flop* model with the same error model as PHYFUM**. The substitution model of this single-sample implementation has 4 parameters, differentiating transitions from homozygous to heterozygous and from heterozygous to homozygous.

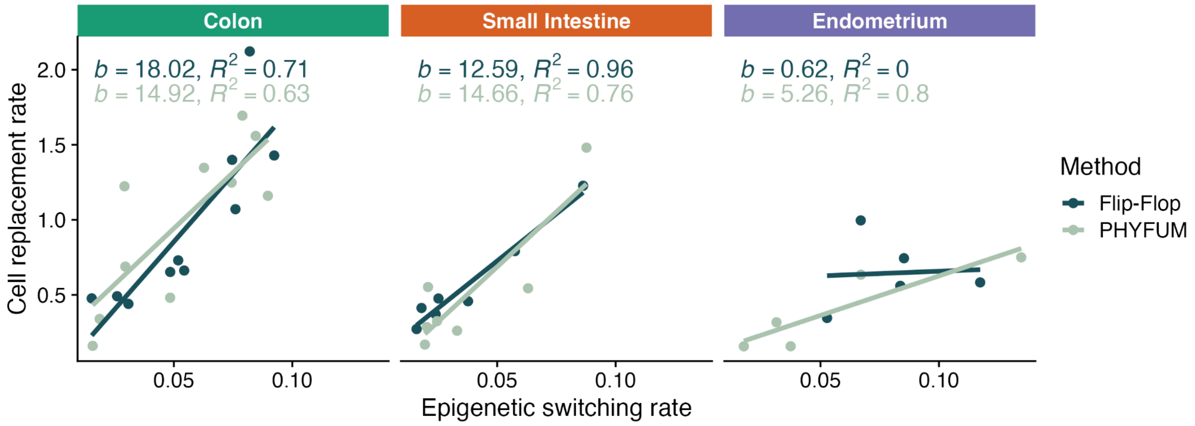


**Supplementary Figure 11: Endometrium-specific decoupling of stem-cell-niche rate estimates in *Flip-Flop* using biological data.** The cell division-induced correlation between clock rates is specifically lost in the endometrium when analysed with *Flip-Flop*, indicating potential parameter identifiability issues in *Flip-Flop*. *Flip-Flop* values correspond to patient means.

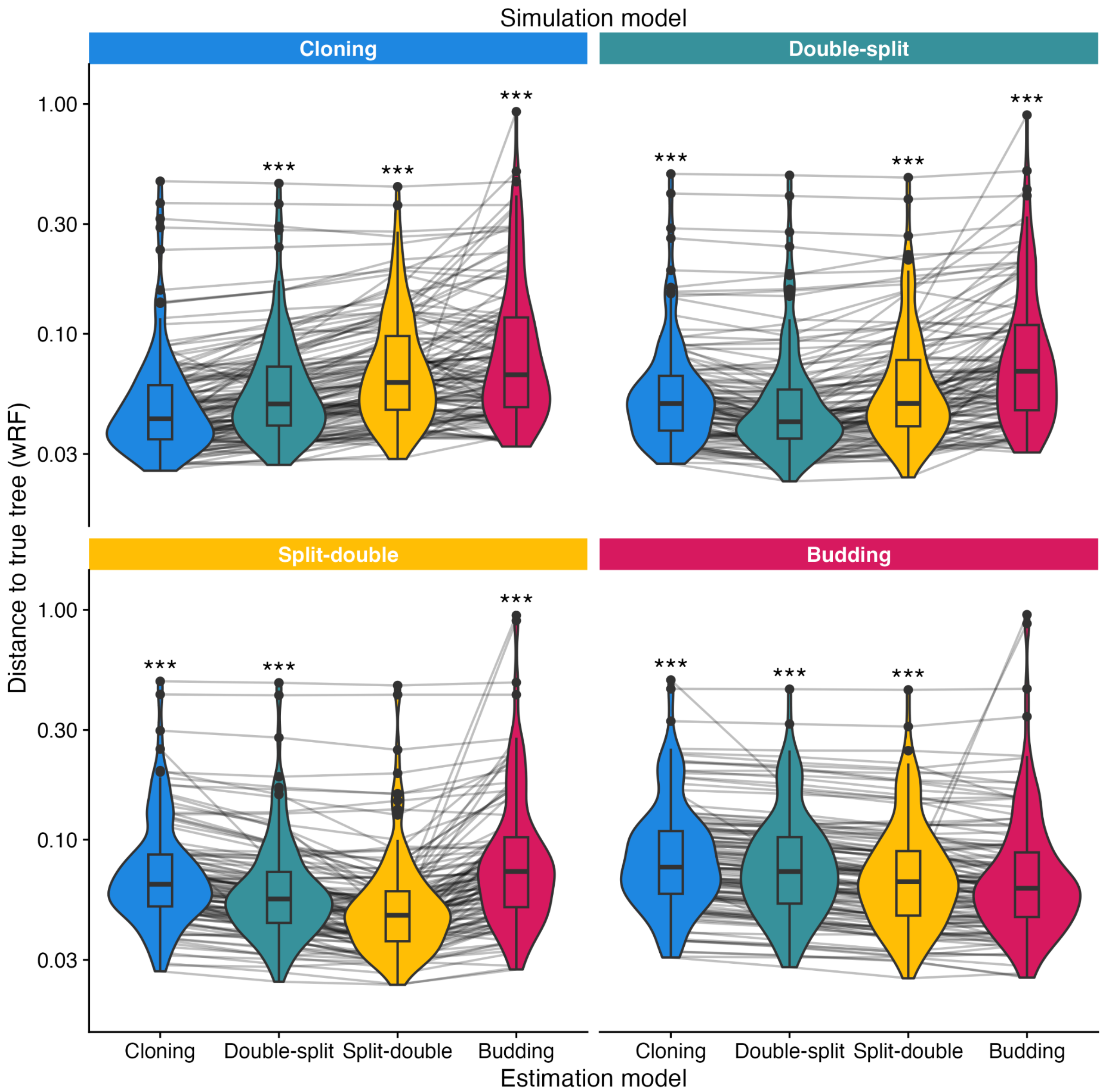


**Supplementary Figure 12**: **Tree estimation error induced by gland-division model mismatch.** Panels depict the true division model of the simulated data. Phylogenetic inference is carried out with all four possible division models for each simulation, and distance to the true simulated tree is calculated with the weighted Robinson-Foulds metric. Paired Wilcoxon signed-rank tests were used to assess statistical differences, with the true model being compared against the other three conditions. Holm-adjusted p-values correcting for all tests in the figure (***: p.adj < 0.001).

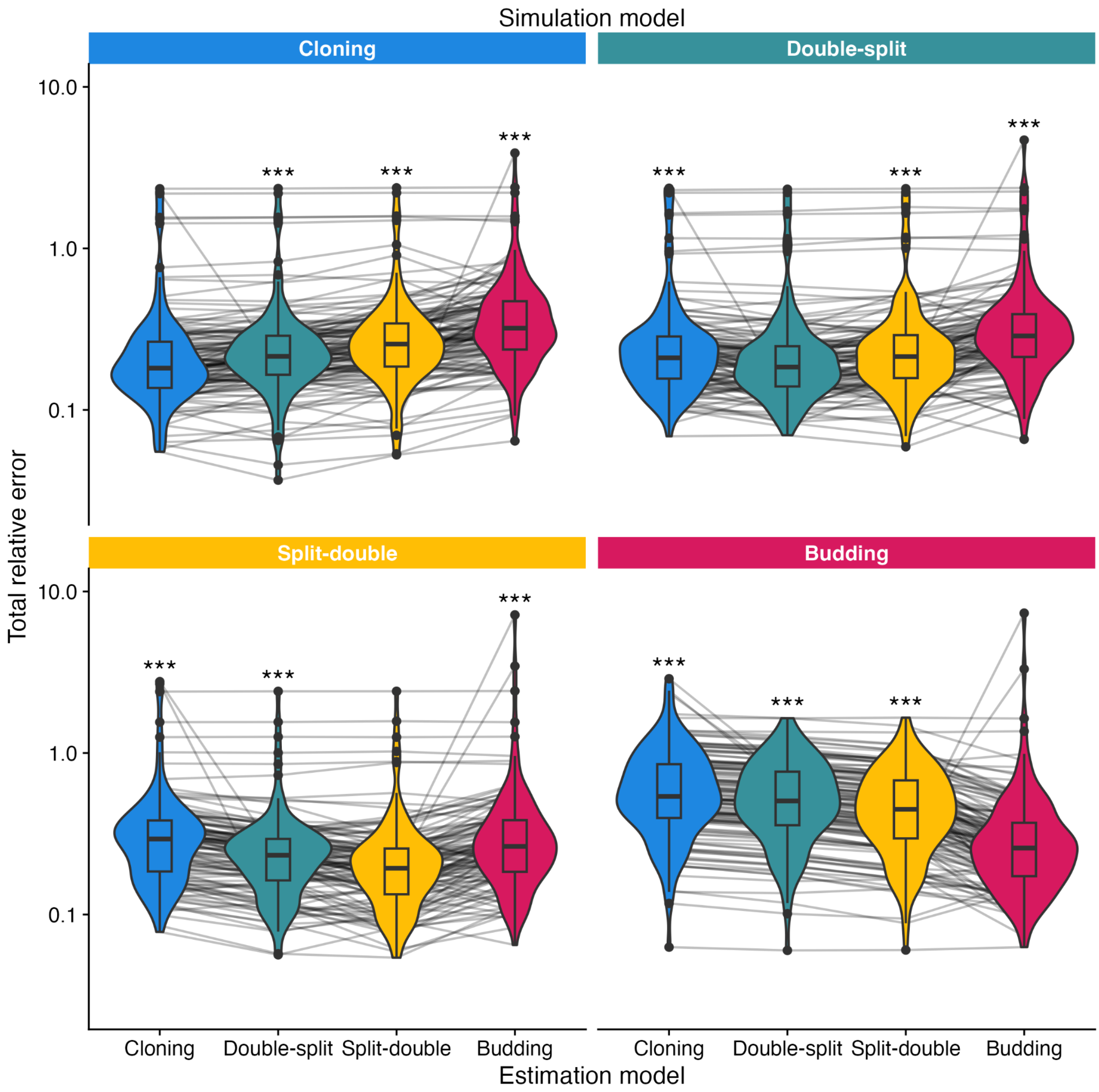


**Supplementary Figure 13: Estimation error induced by gland-division model mismatch.** Panels depict the true division model of the simulated data. The inferred model parameters were compared against simulations' ground truth. The relative error was computed by estimating the absolute difference between each inferred parameter and the ground truth, over the ground truth value. The total relative error comprises the sum of errors of the three fluctuation-methylation model rates and the weighted Robinson-Foulds distance. Paired Wilcoxon signed-rank tests were used to assess statistical differences, with the true model being compared against the other three conditions. Holm-adjusted p-values correcting for all tests in the figure (***: p.adj < 0.001).

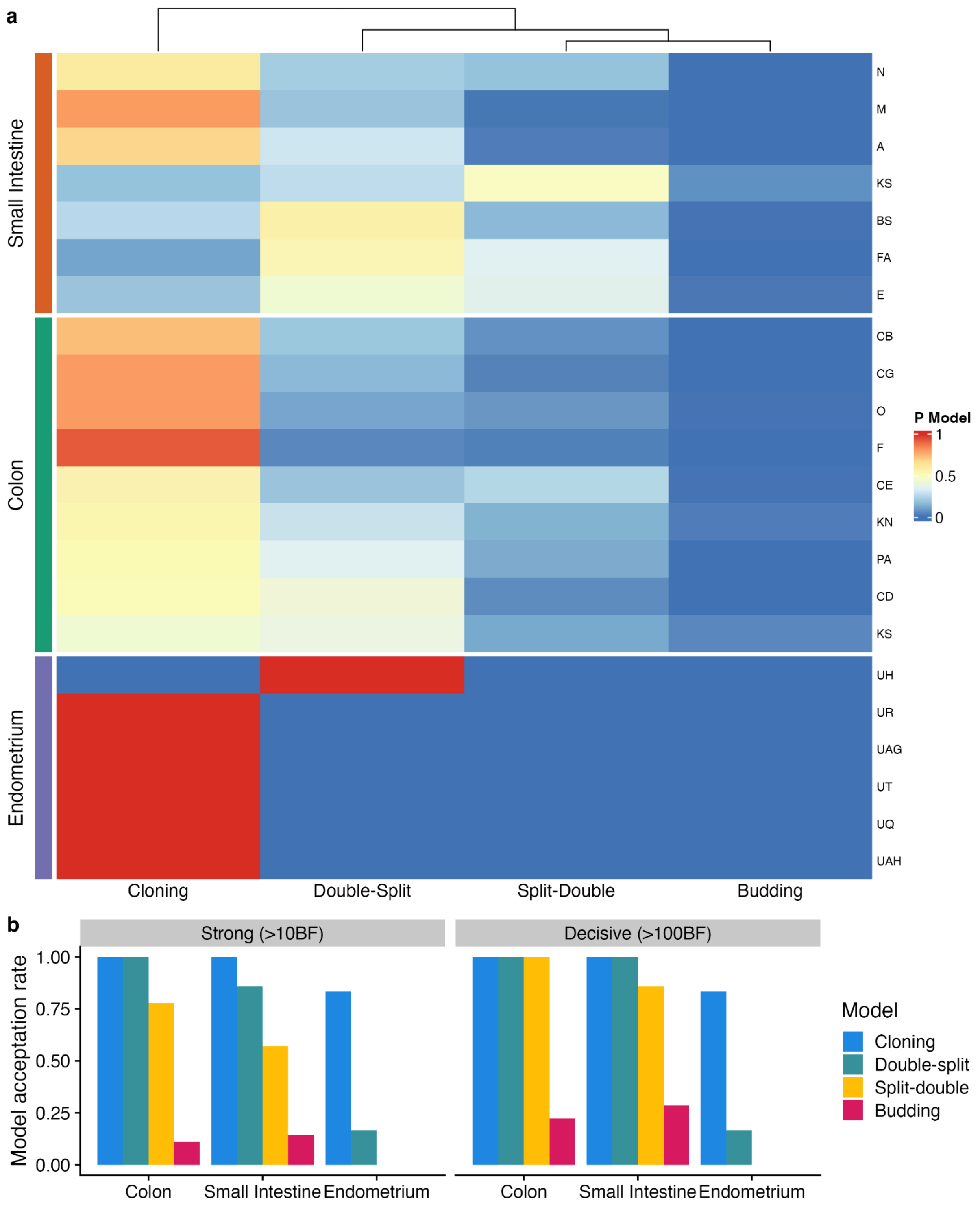


**Supplementary Figure 14: Gland-division model selection in biological data. a,** Posterior probabilities of each gland-division model per patient. **b**, Acceptance rate for each gland-division model on each tissue. The Bayes Factor (BF) is estimated by comparing the best model against the three remaining models. A model is discarded if there is strong (>10BF) or decisive (>100BF) evidence against it when compared to the best one.

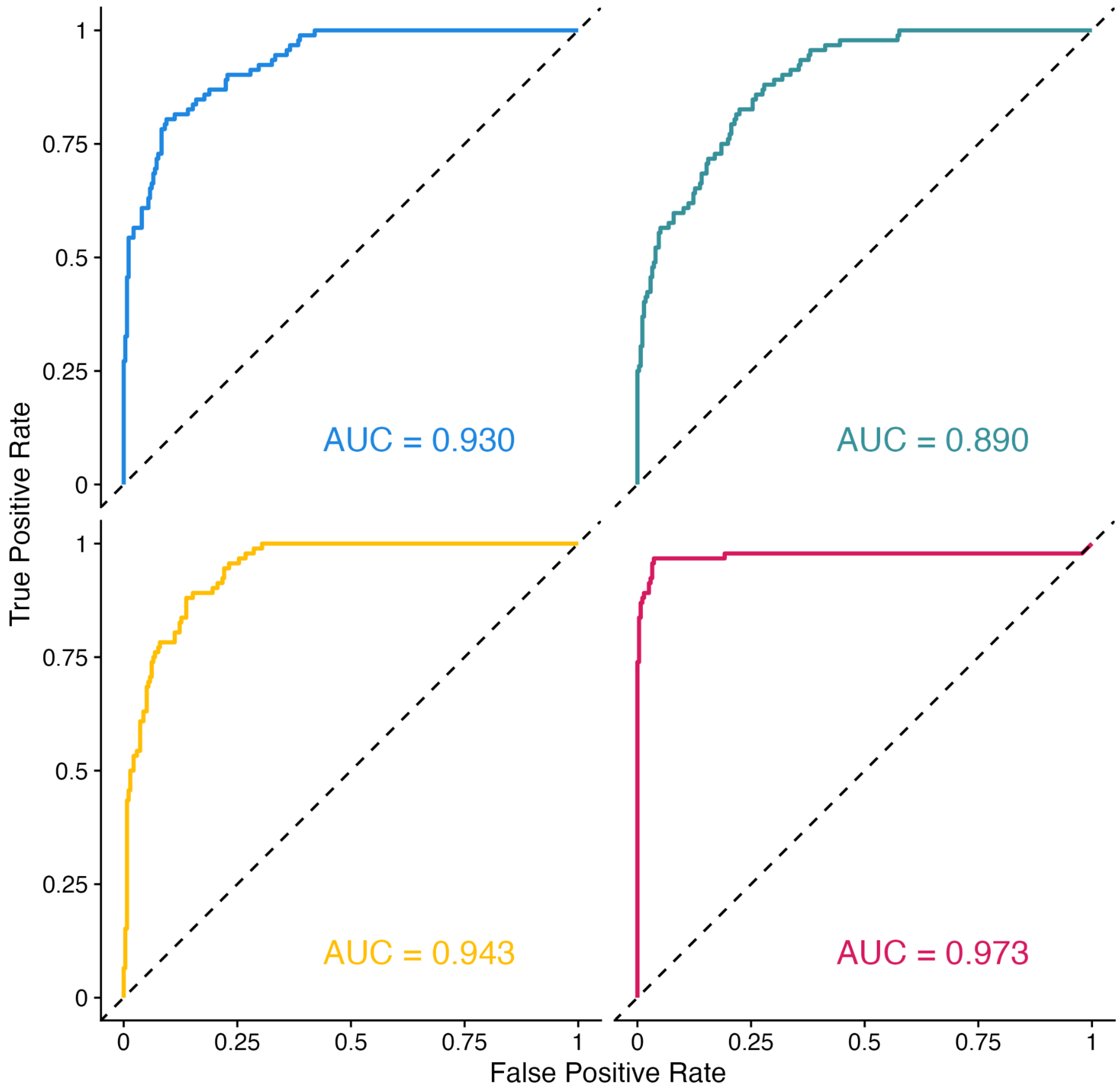


**Supplementary Figure 15: Multiclass ROC curves showing PHYFUM's gland-division classification performance in simulations using the harmonic mean estimator method to estimate the marginal likelihoods.**

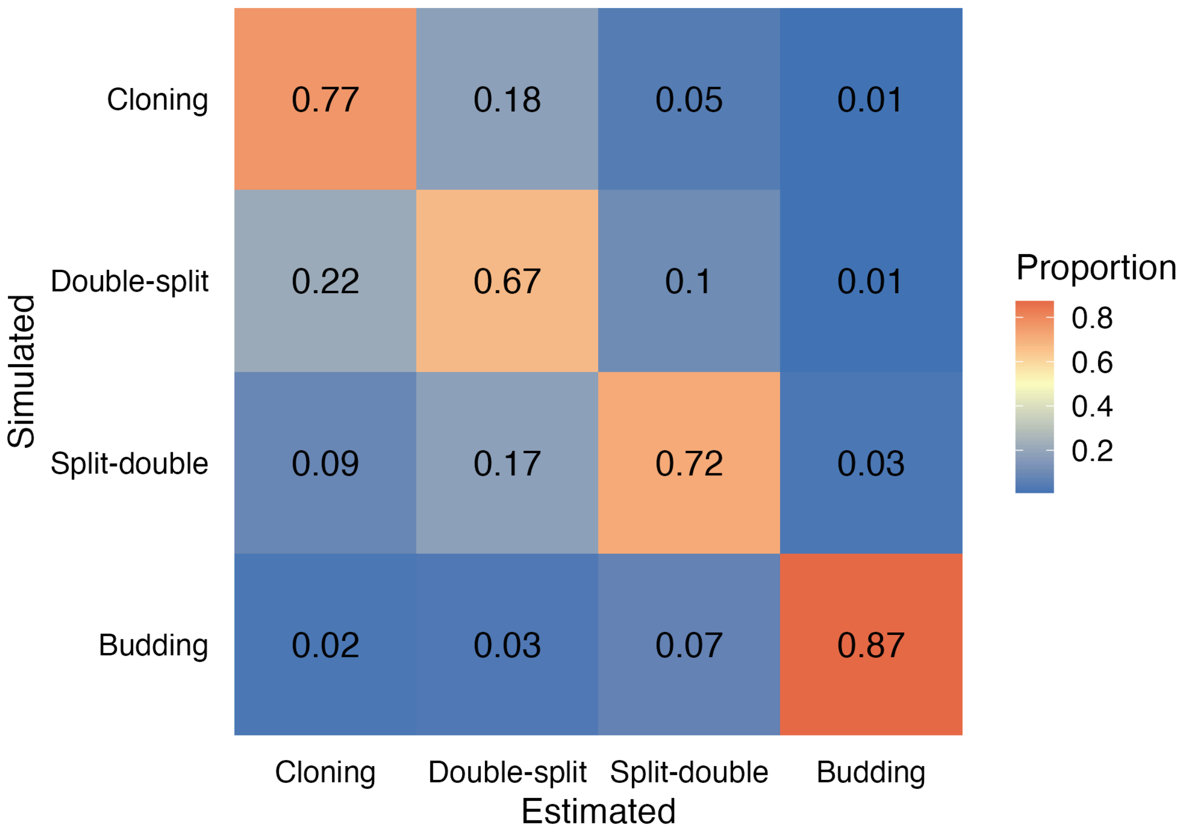


**Supplementary Figure 16: Confusion matrix indicating PHYFUM's gland-division classification performance in simulations using the harmonic mean estimator method to estimate the marginal likelihoods.** Proportion of simulations for which the gland-division model was estimated correctly (diagonal) and incorrectly (off-diagonal).

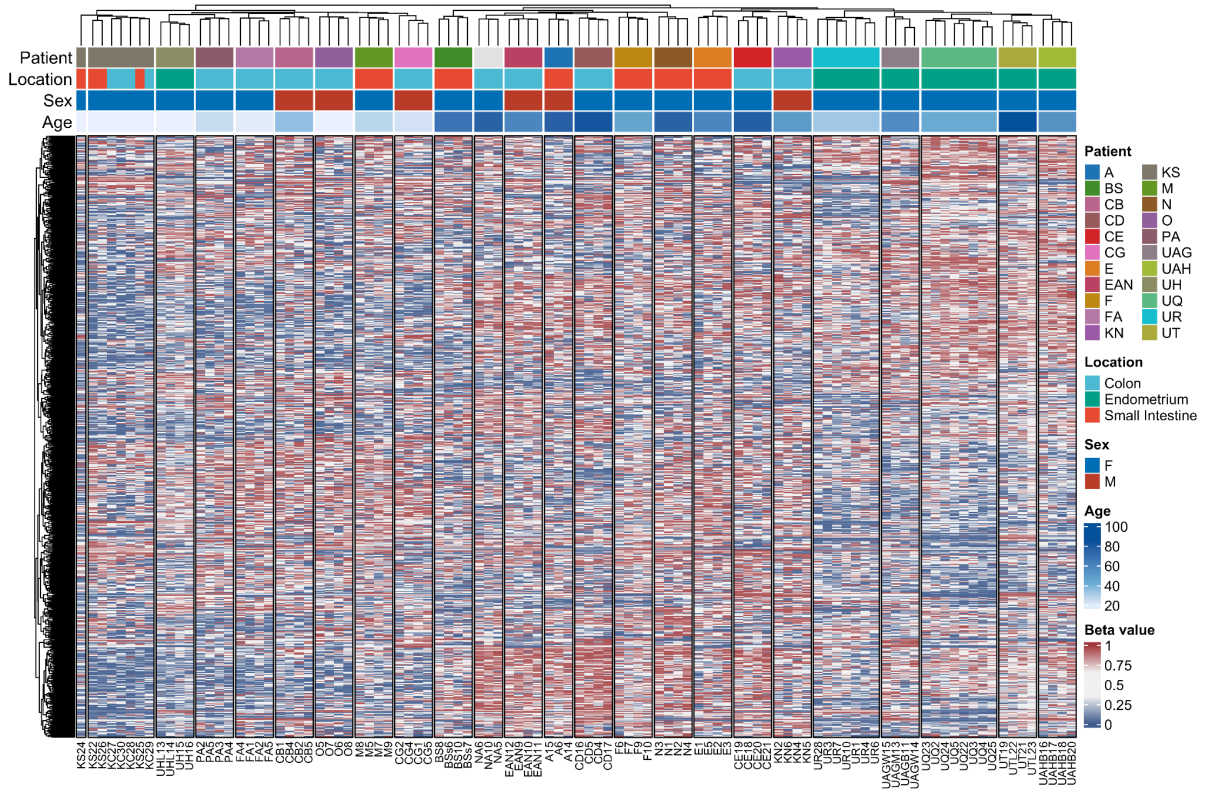


**Supplementary Figure 17: Unsupervised clustering of selected fCpGs in colon and endometrium**. Heatmap of beta values (methylation) of 500 randomly subsampled fCpG sites for plotting convenience.

### Supplementary Tables

#### Suplementary Table S1.  Differences in stem-cell-niche parameter estimates across tissues

| Linear regression | | PHYFUM | | | | | FLIP-FLOP | | | | |
|---|---|---|---|---|---|---|---|---|---|---|---|
| **term** | **variable** | **estimate** | **std.error** | **statistic** | **p.value** | **p.adj** | **estimate** | **std.error** | **statistic** | **p.value** | **p.adj** |
| I(1/age) | gamma | 0.8267 | 0.0364 | 22.6925 | 1.1E-14 | 3.2E-14 | 0.5553 | 0.0790 | 7.0337 | 2.0E-06 | 5.4E-06 |
| locationEndometrium | gamma | 0.0081 | 0.0016 | 5.0008 | 9.3E-05 | 3.7E-04 | 0.0164 | 0.0031 | 5.2450 | 6.6E-05 | 2.6E-04 |
| locationSmall Intestine | gamma | 0.0028 | 0.0015 | 1.8719 | 7.8E-02 | 2.4E-01 | -0.0024 | 0.0031 | -0.7508 | 4.6E-01 | 9.3E-01 |
| I(1/age) | lambda | 20.9543 | 3.3420 | 6.2700 | 6.5E-06 | 6.5E-06 | 18.2569 | 4.1087 | 4.4435 | 3.6E-04 | 3.6E-04 |
| locationEndometrium | lambda | -0.4353 | 0.1479 | -2.9424 | 8.7E-03 | 2.6E-02 | -0.1695 | 0.1624 | -1.0435 | 3.1E-01 | 3.3E-01 |
| locationSmall Intestine | lambda | -0.2715 | 0.1352 | -2.0079 | 6.0E-02 | 2.4E-01 | -0.2057 | 0.1631 | -1.2618 | 2.2E-01 | 9.0E-01 |
| I(1/age) | mu | 0.9002 | 0.0884 | 10.1828 | 6.8E-09 | 1.4E-08 | 0.6692 | 0.0944 | 7.0929 | 1.8E-06 | 5.4E-06 |
| locationEndometrium | mu | 0.0089 | 0.0039 | 2.2692 | 3.6E-02 | 7.2E-02 | 0.0185 | 0.0037 | 4.9608 | 1.2E-04 | 3.6E-04 |
| locationSmall Intestine | mu | -0.0009 | 0.0036 | -0.2488 | 8.1E-01 | 8.1E-01 | -0.0044 | 0.0037 | -1.1874 | 2.5E-01 | 9.0E-01 |
| age | stemCells | 0.0211 | 0.0256 | 0.8247 | 4.2E-01 | 4.2E-01 | 0.0227 | 0.0153 | 1.4816 | 1.6E-01 | 1.6E-01 |
| locationEndometrium | stemCells | -1.5610 | 1.5292 | -1.0208 | 3.2E-01 | 3.2E-01 | -1.2382 | 0.8571 | -1.4447 | 1.7E-01 | 3.3E-01 |
| locationSmall Intestine | stemCells | -1.5623 | 1.3956 | -1.1195 | 2.8E-01 | 5.6E-01 | 0.3047 | 0.8441 | 0.3609 | 7.2E-01 | 9.3E-01 |

| Anova on models | | PHYFUM | | | | | FLIP-FLOP | | | | |
|---|---|---|---|---|---|---|---|---|---|---|---|
| **term** | **variable** | **sumsq** | **df** | **statistic** | **p.value** | **p.adj** | **sumsq** | **df** | **statistic** | **p.value** | **p.adj** |
| age | stemCells | 5.1863 | 1 | 0.6801 | 4.2E-01 | 4.2E-01 | 5.2421 | 1 | 2.1950 | 1.6E-01 | 4.7E-01 |
| I(1/age) | gamma | 0.0043 | 1 | 514.9481 | 1.1E-14 | 3.2E-14 | 0.0016 | 1 | 49.4730 | 2.0E-06 | 1.4E-05 |
| I(1/age) | lambda | 2.7909 | 1 | 39.3125 | 6.5E-06 | 6.5E-06 | 1.6784 | 1 | 19.7449 | 3.6E-04 | 1.4E-03 |
| I(1/age) | mu | 0.0052 | 1 | 103.6892 | 6.8E-09 | 1.4E-08 | 0.0023 | 1 | 50.3090 | 1.8E-06 | 1.4E-05 |
| location | stemCells | 12.6265 | 2 | 0.8279 | 4.5E-01 | 4.5E-01 | 7.3075 | 2 | 1.5300 | 2.4E-01 | 4.9E-01 |
| location | gamma | 0.0002 | 2 | 12.5096 | 3.9E-04 | 1.6E-03 | 0.0012 | 2 | 18.3919 | 5.6E-05 | 3.4E-04 |
| location | lambda | 0.6752 | 2 | 4.7553 | 2.2E-02 | 6.6E-02 | 0.1657 | 2 | 0.9746 | 4.0E-01 | 4.9E-01 |
| location | mu | 0.0003 | 2 | 3.3474 | 5.8E-02 | 1.2E-01 | 0.0016 | 2 | 18.0814 | 6.2E-05 | 3.4E-04 |

### Supplementary Methods

#### 1. PHYFUM: additional implementation details

##### 1.1. Seamless integration of DNA methylation data in BEAST

To address the data type mismatch between the continuous nature of methylation data and BEAST's expected discrete data, we discretized methylation fractions into bins within BEAST. This enabled methylation array data as input to BEAST, which is converted back to float with minimal precision loss prior to inference.

Briefly, let:
P = Precision (number of significant decimal)
$B$ = Number of bins
$S$ = Range start
$E$ = Range end

The number of bins will depend on the precision as:

$$B = 10^{P}$$

The start ($S$) and end ($E$) will depend on the ranges of the variable (0 and 1 in the case of methylation $\beta$ values). Thus, the algorithm creates $B$ bins, representing the space between 0 and 1. The bin width ($\delta$) is calculated then as:

$$\delta = \frac{E-S}{B} = \frac{1}{B}$$

Then, the discretized values, $DV = \{dv1, dv2, ..., dvn\}$ are calculated as:

$$dv_{j} = min(max(\left|\frac{V_{j}-S}{\delta}\right|, 0),\ B - 1)$$

The actual FMC model remaps the values back to float prior to computation, which is done as:

$$v'_{j} = dv_{j} \cdot \delta + S,\ for\ j = 1, 2, ..., n$$

#### 2. PHYFUMr

PHYFUMr (https://github.com/adamallo/PHYFUMr) is an R package that enables users to manually prepare input data and analyze PHYFUM's results, including data preprocessing, copy number variant detection to exclude aneuploid CpG sites, *de novo* identification of fCpG sites, phylogenetic analysis, Bayesian model comparison and/or model averaging for *S* and gland-division model, quality control, summarization, and publication-ready figure generation. Most PHYFUMr functions use pre-existing tools or methods in a convenient fashion for this specific application, but it also introduces novel indices of phylogenetic saturation under the PHYFUM model.

#### 3.1 Measuring phylogenetic saturation

We adapted the index of substitution saturation $I_{SS}$[19], developed for nucleotide sequences, to work with the PHYFUM model. Suppose $N$ samples with $L$ aligned evolutionary characters (i.e., fCpG sites). Each character can take any of the $i = 1,\ldots, C$ categories with a frequency (in all sequences) $P_i$. The index is defined as the ratio of the mean entropy and the expected entropy at full substitution saturation $I_{SS} = \overline{H}/H_{FSS}$ with

$$H_i = -\left(\sum_{j=1}^{C}\left(\frac{N_j}{N}\right)\log_2\left(\frac{N_j}{N}\right)\right),$$

$$H_{\mathrm{FSS}} = -\left(\sum_{N_1=0}^{N}\cdots\sum_{N_C=0}^{N}\left[\frac{N!}{\prod_{i=1}^{C}N_i!}\prod_{i=1}^{C}P_i^{N_i}\sum_{j=1}^{C}\left(\frac{N_j}{N}\right)\log_2\left(\frac{N_j}{N}\right)\right]\right)$$

constrained by $\sum_{j=1}^{C} N_i = N$. When using methylation fractions as input, given PHYFUM's error model parameter values ($\epsilon$, $\Delta$, $\kappa$, and $S$), we assign each methylation fraction to the category that maximizes its likelihood under the error model. With large $S$, the number of compositions to evaluate during $H_{FSS}$ becomes impractically large, and many have negligible effects on the final value due to their low multinomial probabilities. To improve computational efficiency, PHYFUMr offers the option to estimate $H_{FSS}$ by filtering out compositions with multinomial probability under a threshold. For a given dataset, this multinomial probability threshold can be estimated to filter out a desired cumulative multinomial probability by taking $n$ samples ($n$ << compositions of $N$ into $C$ parts) from a multinomial with class probabilities = $\boldsymbol{P}$, calculating their multinomial probabilities, and finding the probability threshold that leaves out the desired cumulative probability.

The $I_{SS}$ index is oblivious to phylogenetic saturation in the LCA-MRCA branch. In order to complement this index, PHYFUMr also calculates the Earth Mover's Distance between the prior state frequencies at the LCA and the state frequencies $\boldsymbol{P}$ relative to the maximum distance to any $\boldsymbol{P}$.

### 3. PHYFUMflow

To facilitate the analysis of fluctuating methylation clocks, we developed a Snakemake workflow, PHYFUMflow, that runs an end-to-end PHYFUM analysis. Users can provide either raw methylation data (IDAT files) or pre-processed beta values.

When raw files are supplied, the pipeline utilises *minfi*[20] with FunNorm normalisation to generate methylation beta values. Next, if desired, the pipeline automatically detects and removes CpG sites suspected to lie within regions affected by copy-number alterations. This is accomplished using *Conumee*[21] and *rascal*[22]. Segments exhibiting absolute copy number greater than ±0.25 (after purity correction) compared to the baseline are classified as altered, and CpG sites within these segments are blacklisted and not used during fCpG selection. Alternatively, users can supply a custom blacklist of CpG sites.

For datasets comprising multiple individuals, PHYFUMflow can infer FMCs *de novo*, as described in this manuscript, while considering the new fCpG phylogenetic saturation statistics developed here and implemented in PHYFUMr. Otherwise, the user must provide a pre-specified list of CpG sites. Next, PHYFUM is executed in parallel across multiple independent MCMC chains for all specified values of $S$ and gland-division models per patient. The conditional model evidence is estimated for each using either the harmonic mean estimator (HME), the stepping stone method, or the path sampling method. Finally, PHYFUMflow uses PHYFUMr to conduct Bayesian model comparison and/or model averaging to either select the optimal *S* and provide conditional parameter estimates, or marginalize over *S* to provide unconditional estimates, respectively. The workflow also performs automated downstream plotting and quality-control analyses using the multiple MCMC chains per condition to ensure mixing and convergence of continuous parameters and tree topology.

PHYFUMflow (https://github.com/pbousquets/phyfumflow) is distributed as a Python package and as a Docker image bundled with PHYFUM, providing a streamlined solution for researchers studying clonal dynamics in healthy and pre-cancerous tissues.

#### 4. PHYFUM simulator

We designed a simulator under the PHYFUM model to validate PHYFUM's implementation and study its accuracy. Users can provide a tree describing the evolution of several stem-cell niches or let the program simulate a coalescent tree for a specified period of time with msprime[23].

The program initialises a set of CpG sites, either fully methylated or unmethylated, for all $S$ stem cells. Afterwards, the fCpG sites evolve according to a predefined transition matrix, until the tree-branch splits. The states are stored, and the sites are again let drift independently for each new daughter branch until a new bifurcation occurs. This is done successively until the tree tips are reached. Afterwards, noise is added following the error model.

At each internal node, the program can use the different stem-cell-niche division models implemented in PHYFUM. By default, the internal node states are directly inherited by the daughter lineages as the starting states of their simulation (Cloning model). Alternatively, all stem cells may duplicate and then distribute at random between their daughters (Double-Split model), unevenly distribute the stem cells and then duplicate (Split-Double), or, more extremely, a single stem cell leaves the ancestral stem-cell niche to create a new one, and the rest remain together (Budding). Importantly, the Budding simulation model departs slightly from the Budding estimation model, since the estimation model assumes the ancestral stem-cell niche remains unchanged, while the simulation model assumes the stem cell that left is replaced by one of the existing ones. Users may provide the individual age, the number of fluctuating CpG loci to simulate, the model parameters and hyperparameters, and the gland-division model. Alternatively, the user can let the program select plausible random values for the parameters and hyperparameters.

### Supplementary Note 1

#### Effect of marginal likelihood estimation method in Bayesian model comparison for stem-cell number estimation

Bayesian model comparison is typically computationally expensive, as it requires sampling from a series of power posteriors to estimate each marginal likelihood using stepping stone[14] (SS) or path sampling[15](PS). Alternatively, the harmonic mean estimator (HME) only requires samples from the posterior, but it can fail catastrophically when its variance becomes very large. We investigated the relative performance of HME, SS, and PS in estimating PHYFUM's *S* (number of stem-cells in the stem-cell niche) via Bayesian model comparison using both our biological data and a simulation study.

Using the biological data from the main manuscript but a different set of fCpG sites (this study was carried out during the first stages of software development), we compared the best-fit *S* values across the cohort (Supplementary Note 1 Figure 1) for the three marginal-likelihood estimation methods. HME overestimated the marginal likelihoods compared with SS and PS, but showed very similar relative values across *S* per patient, resulting in adequate model-selection accuracy: only 4/24 cases had errors larger than 1 stem-cell, and 11/24 cases had identical best-fit solutions.

To actually quantify the differences in estimation accuracy, we conducted a simulation study for which we generated 36 simulations with 2,000 fluctuating CpG sites and all combinations of several parameter values:

- Stem cells, $S$: [4, 7, 10]
- Methylation rate, $\mu$: [0.005, 0.01]
- Number of samples, *N*: [5,9]
- Expected number of fixations per site due to cell-replacement, *r*: $[r_1,r_2,r_3] \sim N(10,5)$, sampled per combination of the previous 3 parameters

- Stem cell replacement rate, $\lambda$: $r(S - 1)^2/(tS)$
- Demethylation rate, $\gamma \sim \mu N(1,0.04)$
- Phylogenetic tree: *N*-tip coalescent sampled per condition

- Assay precision, $\kappa = 100$
- Sample age, $t = 45$
- Maximum observable methylation fraction $\beta$, $\epsilon = 0.92$
- Minimum observable methylation fraction $\beta$, $\Delta = 0.04$
- Gland-division model: *Cloning*

We ran two independent 750k-iteration MCMC estimation chains per condition, sampling every 75, and discarded the first 10% of each as burn-in, followed by marginal likelihood estimation for each (sampling 100 power posteriors with the same total number of iterations as the MCMC, with powers ~ beta(0.3,1)). All other PHYFUM parameters were set as default in PHYFUMflow unless specified. We ran the estimation for each condition with 10 different stem niche sizes (estimation S, *eS*) [3,4,5,6,7,8,9,10,11,12], yielding a total of 360 estimation conditions.

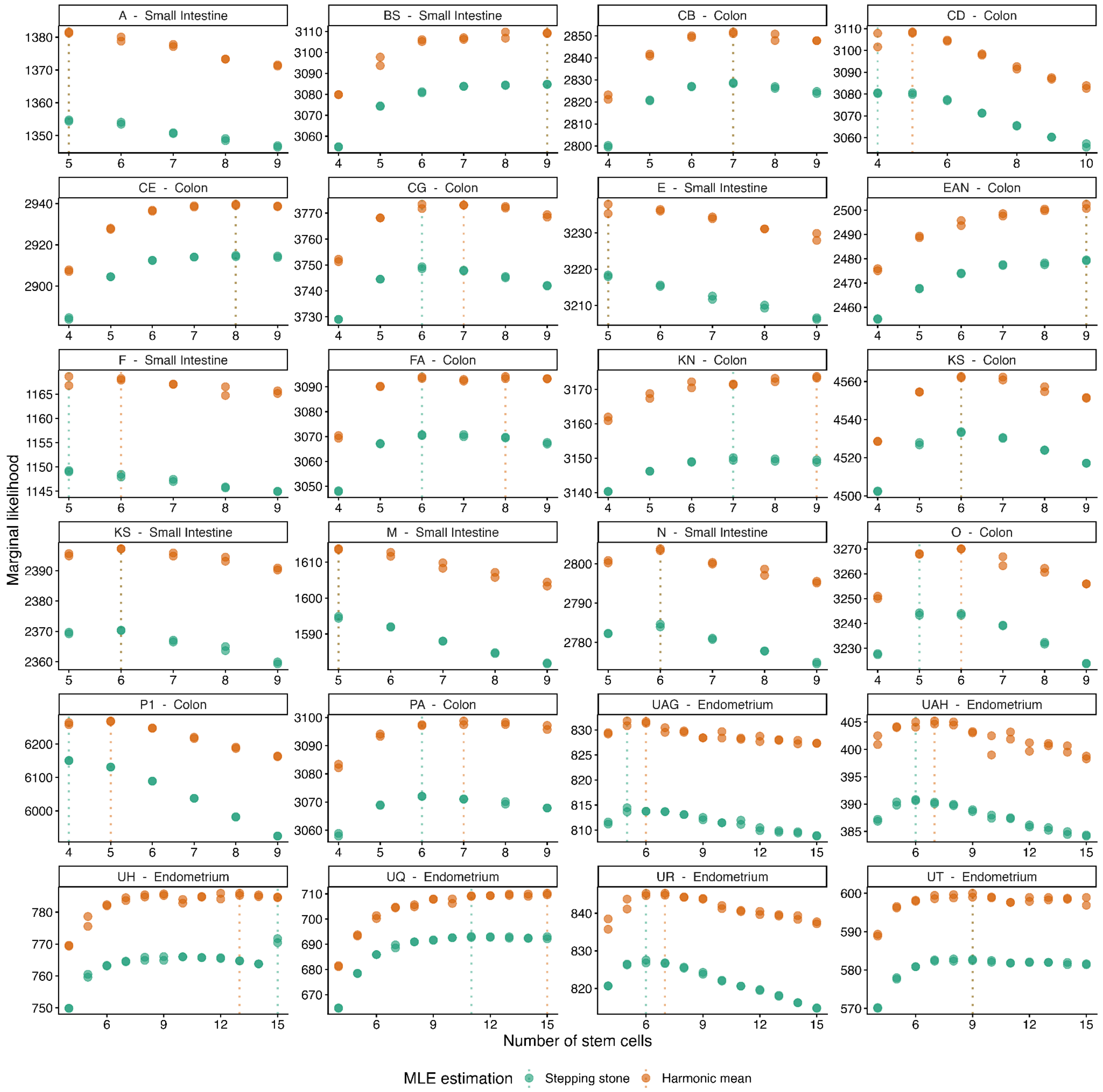


**Supplementary Note 1 Figure 1**: **Comparison of Harmonic Mean Estimation and Stepping Stone methods for model comparison of the *S* parameter on real data**. Vertical dotted lines depict the optimal solution for each method. These results were obtained during development using a different set of fCpG sites and therefore do not exactly match those presented in the main manuscript. Path Sampling results are identical to Stepping Stone results with regard to the best-fit *S* values, and they are not plotted for simplicity.

We calculated the estimation accuracy of the three marginal likelihood estimation methods for both the point-estimate best-fit *S* and taking into account reconstruction uncertainty by considering any *S* within 10 Bayes factors of the best (i.e., rejecting *S* values with strong evidence against them) (Supplementary Note 1 Figure 2). These results show that HME accuracy is comparable to that of SS and PS in this application. Point-estimate accuracy decreases with *S* for all methods, but the true model is contained within the set of accepted *S* values with high accuracy across *S* for all methods.

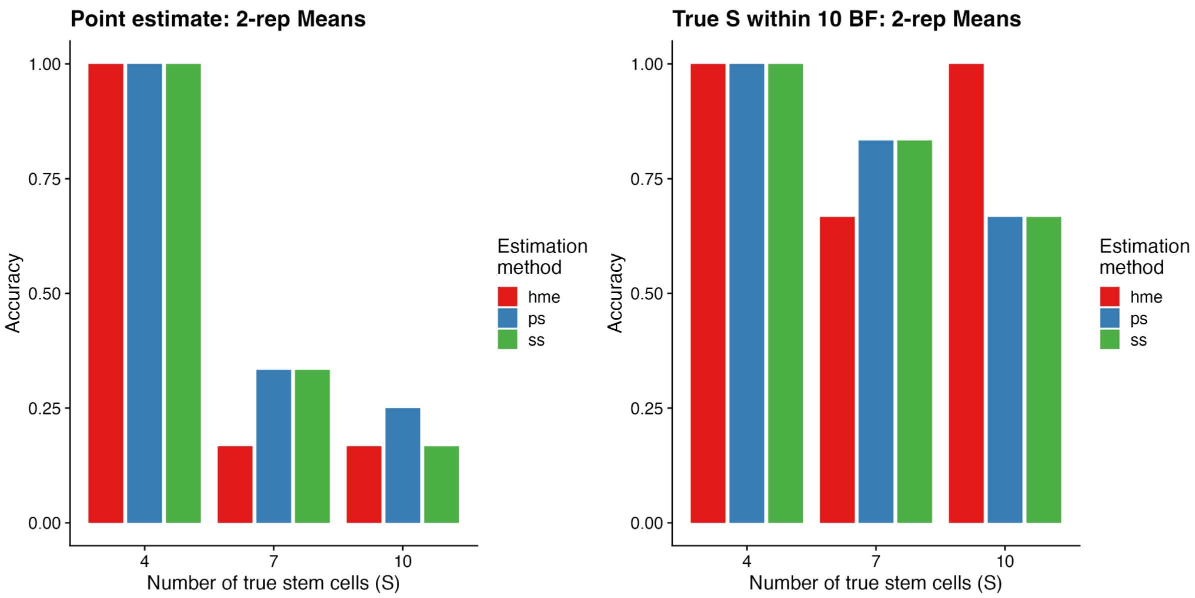


**Supplementary Note 1 Figure 2: Comparison between model selection strategies in the accuracy of selecting *S* using simulated data.** Left: point-estimate accuracy, Right: accuracy of the true model being contained within 10 Bayes Factors of the best model. Hme: harmonic mean estimator, ps: Path Sampling, ss: Stepping Stone.
